# Fungal Pathogen Gene Selection for Predicting the Onset of Infection Using a Multi-Stage Machine Learning Approach

**DOI:** 10.1101/2023.09.26.559518

**Authors:** Graham Thomas, Oliver Stoner, Fabrizio Costa, Ryan M Ames

## Abstract

Phytopathogenic fungi pose a serious threat to global food security. Next-generation sequencing technologies, such as transcriptomics, are increasingly used to profile infection, assess environmental adaptation and gauge host-responses. The accumulation of these large-scale data has created the opportunity to employ new computational methods to gain greater biological insights. Machine learning approaches, that learn to identify patterns in complex data sets, have recently been applied to the field of plant-pathogen interactions. Here, we apply a machine learning approach to transcriptomics data for the fungal pathogen *Zymoseptoria tritici*, to predict the onset of infection as measured by timing of the appearance of necrosis. We present a method for identifying the most important genes that predict infection timings, accurately classify isolates as early and late infectors and predict the timing of infection of ‘novel’ isolates using only a subset of the data. These methods and the genes identified further demonstrate the use of these tools in the field of plant-pathogen interactions and have implications for the identification of biomarkers for disease monitoring and forecasting.

Fungi that infect plants pose a serious threat to global food security. Methods to study these pathogens generate vast amounts of data that create new opportunities for computational tools to analyse them. Machine learning methods can learn patterns in complex data such as when genes are turned on or off in fungal plant pathogens. In this study we use machine learning approaches to predict the onset of infection in several isolates of an important fungal pathogen. We show that these methods can identify a small group of genes that are predictive of the infection onset. We can even use these methods on ‘novel’ isolates to infer the likely timing of disease development. Our work has implications for plant disease diagnosis, monitoring and forecasting.

## Introduction

Phytopathogenic fungi threaten global food security, destroying up to 30% of crop products through disease and spoilage [1, 2]. To cause disease and complete their life cycle, plant pathogens have to overcome the physical and chemical barriers of plant immunity, including pathogen-associated-molecular-pattern (PAMP)-triggered-immunity (PTI) [3]. They achieve this by secreting effector proteins, which disrupt host signalling for threat perception and defence mechanisms [3–5]. In turn, plants have evolved resistance (R) proteins which detect specific pathogen effectors or monitor the effector-targeted proteins and initialise effector-triggered immunity (ETI) [6]. These gene-for-gene (R gene to effector gene) interactions can determine the success or failure of infection [7]. ETI induces a suite of responses including a hypersensitive response (HR), localised cell death, activating multiple protein kinase cascades, as well as initiating an oxidative burst and promoting expression of many defence related genes [3, 8].

The Dothidiomycete fungus *Zymoseptoria tritici* is the causal agent of Septoria tritici blotch (STB), a severe disease of wheat [9]. STB results in significant economic losses through reduced yields of 5-10% per annum in the EU [10] and the cost of disease management through application of fungicides, which accounts for approximately 70% of the European agricultural fungicide market [11, 12]. *Z. tritici* is capable of rapid evolution due to a high rate of gene flow and recombination in large populations [13–16]. This fact, coupled with management practices has driven selection in *Z. tritici* for resistance to all major fungicides [17–23]. The situation is compounded by *Z. tritici’s* highly plastic genome, which enables the fungus to overcome major wheat resistance genes [24]. *Z. tritici* has 21 chromosomes with eight chromosomes being dispensable and absent in some isolates of the species [25]. Indeed, 40% of the *Z. tritici* pan-genome, the entire set of genes within the species, is composed of orphan genes [26, 27], giving *Z. tritici* the largest known accessory genome among fungi [28]. Deletion of the dispensable chromosomes has confirmed that some are necessary for host specificity [29]. Transcriptomic studies show flexibility of infection programs with heterogeneity in gene expression throughout development between isolates [30, 31]. This extends to variability in development and aggressiveness when assessing specific isolate pathogen-host combinations [32].

*Z. tritici* is a member of the Mycosphaerellaceae family, which are often deemed hemibiotrophic due to having a long asymptomatic, biotrophic phase in the disease cycle [33]. This is followed by a rapid switch to a necrotrophic phase and the development of disease symptoms, including irregular chlorotic lesions and then necrotic blotches forming on the leaf [34–36]. Necrotrophy is associated with the hallmarks of PTI and ETI in the host, including programmed cell death (PCD) and accumulation of H_2_O_2_ [37, 38]. This results in the collapse of mesophyll tissue and an abundance of nutrients flooding the apoplastic spaces, facilitating rapid fungal growth and finally sporulation [39–42]. With limited evidence of biotrophy during the asymptomatic phase, *Z. tritici* is more accurately described as a latent necrotroph [43]. The asexual reproductive cycle is characterised by distinct developmental stages throughout both the asymptomatic and necrotrophic phases, lasting 2-3 weeks [30, 44, 45]. A spore will alight upon a wheat leaf, develop invasive hyphae, gain ingress through natural openings (stomata) as opposed to forming specialised structures such as an appressorium, and growth is exclusively in the intercellular spaces between the mesophyll cells of the leaf. These stages comprise the asymptomatic phase of infection and typically last 914 days [35, 44, 46–50]. The necrotrophic phase is represented by increased fungal biomass, hyphae reaching new stomata and forming new fruiting bodies called pycnidia in the sub-stomatal cavity. Mature pycnidia are packed with macropycnidiospores, which are released through the stomatal aperture under conditions of high humidity and spread by rain splash to repeat the infection cycle [11, 50, 51]. Development is asynchronous between individual spores infecting a single leaf [45] and spores can grow epiphytically in excess of 10 days [52].

Although the stages of development are known, a complete picture for the molecular mechanisms controlling development remain to be established. Efforts to elucidate this subject have focused on characterising virulence factors, focusing on small secreted effectors [42, 44, 53–68], transcription factors and kinases [69– 74]. A broader approach involves transcriptome profiling through the infection cycle, assessing systems-level adaptation to environmental changes and differences in host responses from cultivars with ranging susceptibilities [30, 31, 44, 75–81]. Transcriptome studies generate vast amounts of data and we require intelligent methods to mine and understand them. Recently, machine learning approaches have been applied to a range of areas in plant-pathogen interactions including disease monitoring, genomic selection for resistance and the identification of potential effectors (reviewed in [82]). Although studies have looked at using a variety of transcriptomic data to identify stress conditions [83] and genes associated with complex agronomic traits in host plants [84], to the best of our knowledge no work has focused on using pathogen transcriptomic data to predict the outcome of infection and identify the key drivers, or genes that predict these outcomes.

In this work, we apply a combined Machine Learning (ML) approach with a parametric statistical model to available *Z. tritici* transcriptomic data captured during infection to: (i) identify a small subset of genes which may be associated with infection progression; (ii) uncover relationships between expression changes of these genes over time and the timings of necrosis appearance as a marker of infection outcome; (iii) predict how quickly a ‘novel’ pathogenic isolate reaches this infection milestone based on the expression of the subset of genes. Our approach relies on little to no expert information about novel isolates and is generally applicable across different experimental designs (e.g. in the timings of observations) and species. The method we propose is composed of three stages: in the first stage we use a ML approach to identify the genes which best characterise the infection phases; in the second stage, we quantify how the expression of these genes changes over time; and in the third stage, we develop a statistical model to give a “high resolution” estimate of the timing of the infection “peak”. The key insight in stage one is to formulate the problem as a feature selection problem in a binary classification task, where the two classes/conditions to discriminate are the non-infectious state and the infectious state. The key insight in stage two is the definition of a robust notion of distance from the non-infectious state, to this end we employ the Mahalanobis distance between gene expressions (so to take into account the natural variability of each gene) but limiting the information to the (few) discriminative genes identified in stage one (so to achieve the largest possible signal to noise ratio). Finally, the key insight in stage three is to use the metric developed in stage two to quantitatively define specific phases in the shape of the temporal evolution of the infection, namely: an onset phase, a peak phase and a decay phase to a stable condition.

The empirical results show that our approach can identify a characteristic subset of genes that significantly change expression between the onset of infection and appearance of necrosis. Furthermore, summarising the change in expression of these genes is sufficient to differentiate between isolates showing an early or late appearance of symptoms. Finally, we show that our methods are potentially capable of predicting infection timings of ‘novel’ isolates, even on a subset of the available transcriptomics data. Overall, our results suggest potential utility of the proposed approach for the identification of a small subset of genes predictive of an infection outcome that can be investigated as potential candidates for disrupting the disease process as well as for forecasting the emergence of new plant disease.

## Results

### Data acquisition and experimental design

We used gene expression data from two studies; *(i)* Haueisen *et al*. characterised gene expression for 3 isolates of *Z. tritici* (Zt05, Zt09 and Zt10) at 4 stages of infection (A, B, C and D) determined by confocal microscopy [30]. Zt05 is a field isolate from Denmark, Zt09 is a derivative of the reference isolate IPO323 and Zt10 is another field isolate from Iran. Previous work has identified *>*500,000 and *>*600,000 single nucleotide polymorphisms (SNPs) with reference isolate IPO323 indicating there is significant genetic distance between these 3 isolates [85]. *(ii)* Palma-Guerrero *et al*. recorded gene expression data for 4 isolates of *Z. tritici* (3D7, 3D1, 1A5 and 1E4) at 7, 12, 14 and 28 days post infection (dpi) [31]. These isolates were all collected from two wheat fields in Switzerland, but show considerable genetic diversity with approximately 310,000 SNPs between each pair [75, 79]. Selection of isolates from diverse geographical regions that show significant genetic diversity allows our approach to be generalised across *Z. tritici* isolates. Normalising and filtering the data (see Materials and Methods) left 9,371 and 6,641 genes in the Haueisen *et al*. and Palma-Guerrero *et al*. data respectively, with 6,486 genes present in both data sets. Only genes detected to be expressed in all isolates were used in this study, this is important as identifying commonly expressed genes that are predictive of infection allows us to demonstrate the utility of these methods across isolates.

Both sources of data for this study report a variety of infection outcomes including the timing of disease symptom development and the percentage leaf area covered by necrosis and pycnidia. For this study we used the most comparable measure common to both experiments, which was the timing of the appearance of necrosis measured in days post infection (dpi), which we used to classify infection by these isolates as early or late onset. This follows the findings that isolate Zt10 develops necrosis significantly later than isolates Zt05 and Zt09 [30], and that symptoms of infection, including necrosis, were also found to occur later and be less severe for isolate 3D1 compared to 3D7, 1A5 and 1E4 [31].

### Machine learning to identify genes predictive of infection timings

We first used a random forest classifier to identify those genes that change their expression significantly between two distinct time points in the infection period that therefore, may represent important genes in disease progression. For the first time point, we selected the initial time point of infection. For the second, we selected the time point closest to necrosis development, the infection outcome of interest. The time points closest to necrosis development were infection stage C and 14 dpi for the Haueisen *et al*. [30] and Palma-Guerrero *et al*. [31] data, respectively. These two time points are the two “classes” the random forest seeks to distinguish between based on the gene expression values, which are the “features”. This setup allows us to quantitatively assess how predictive each gene is in determining at what stage of infection the observation is made. The random forest classifier had an out-of-bag prediction error of 3.5%, which is the fraction of miss-classified samples. The 20 most important genes for determining the stage of infection are reported in Table 1, as quantified by their impurity score from the random forest classifier.

**Table 1.**
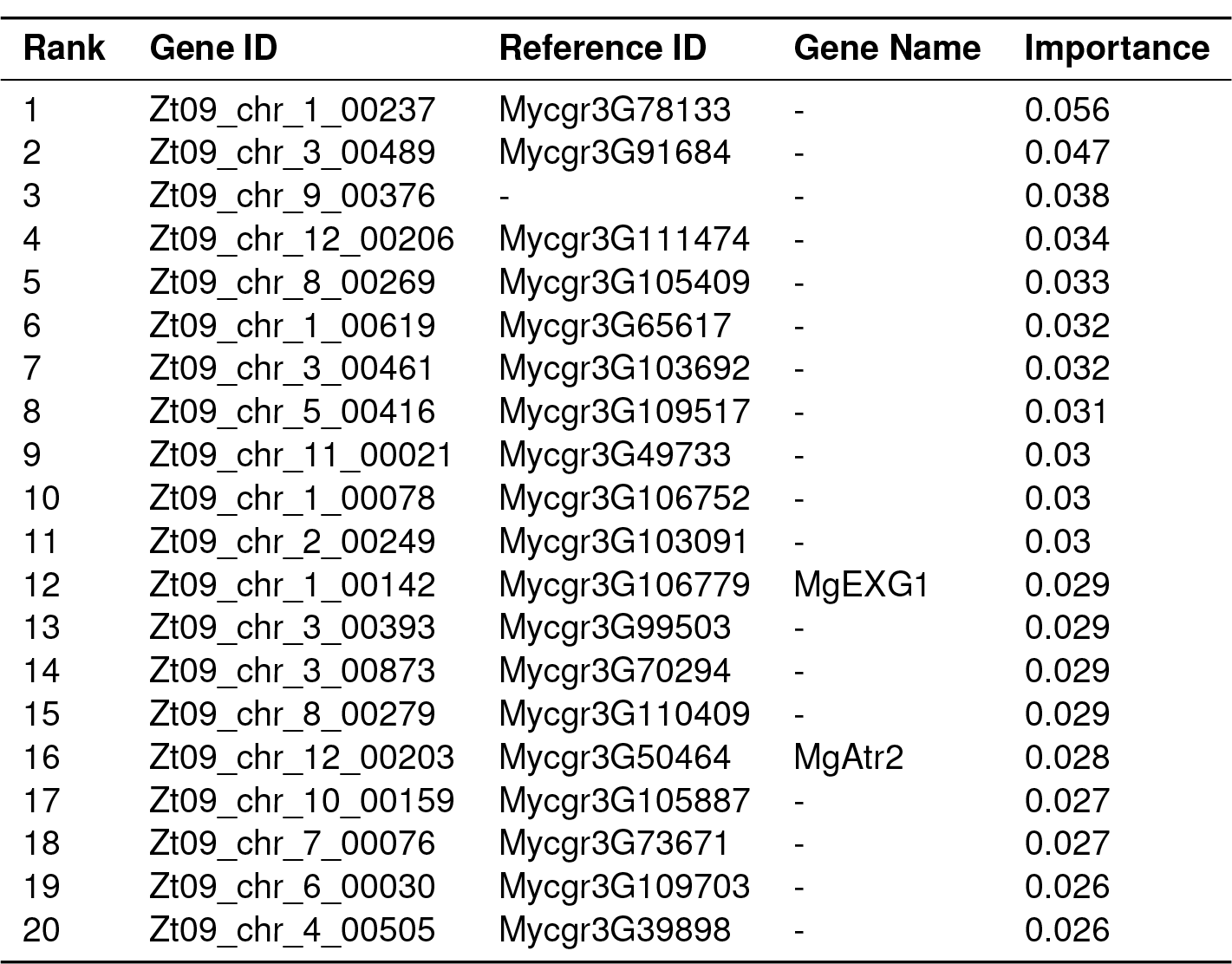
The 20 most important genes as quantified by their (rounded) impurity scores in order of decreasing rank, with all isolates as inputs.

### Comparing machine-learning to differential expression analysis

Traditionally, to identify genes that change in their expression significantly over time, we would use standard differential expression analysis rather than the machine learning approach described above. The rational for using a ML feature selection approach is that, due to the complexity of the interactions in biological systems, we should strive for methods that can process correlation and non linear interaction effects rather than limiting the selection process to univariate approaches based on t-tests – or other statistical tests – that model one single gene at a time [86, 87]. To compare our results to those of the more traditional analysis, we identified significantly differentially expressed genes using edgeR [88]. This analysis revealed 2,757 significantly differentially expressed genes (*corrected p <* 0.05), of which, all 20 genes identified in Table 1 are present (12 up-regulated, 8 down-regulated; full list available in Table S1). However, our most important genes are not the most significant according to *p −value* ranking, nor do they show the largest changes in expression ranked by fold change. In our approach these genes have been selected on the basis of their joint capacity to inform the classifier to distinguish the two conditions, irrespective of the magnitude of their expression levels.

It is not uncommon for conventional differential expression analysis to reveal very large lists of genes and, without significant further analysis, including arbitrary cutoffs, it would be difficult to narrow these lists down to a manageable number of genes for further analysis and potential experimental investigation. We believe that machine learning based methods for gene selection offer an attractive alternative to traditional differential expression as they 1) allow to rank genes using a multivariate importance measure (e.g. the reduction of the classifier discriminative power) and they often 2) yield sorted lists of genes’ importance scores with a pronounced “knee”, which allows the identification of a natural cutoff.

Haueisen *et al*. have also published lists of differentially expressed genes that represent a core transcriptional program during wheat infection [30]. These genes are differentially expressed between infection stages across all 3 isolates present in the study. In total the authors identify 676 genes that show differential expression between the 4 identified stages of infection. As mentioned above, this represents a relatively large number of genes to process. In this study, 8 of the 20 most important genes are also differentially expressed between stages of infection in Haueisen *et al*.. All of these 8 genes (Zt09_chr_1_00237, Zt09_chr_3_- 00489, Zt09_chr_9_00376, Zt09_chr_3_00461, Zt09_- chr_2_00249, Zt09_chr_12_00203, Zt09_chr_6_00030 and Zt09_chr_4_00505) show differential expression at stage C (note only consecutive stages were tested i.e. B vs C and C vs D), and in all cases except Zt09_- chr_4_00505, these genes are up-regulated at stage C. This is an expected result as stage C relates to necrosis development, which is our infection outcome of interest. This is the same time point that we have used to identify the most important genes presented in Table 1. Although there is considerable overlap between the most important genes identified in this study and those genes showing differential expression in the core transcriptional response identified by Haueisen *et al*. [30], there are 12 genes that do not overlap. This may be because our study uses a wider range of isolates with more diversity, potentially allowing us to capture a more general set of genes that define infection progression.

### Summarising change of expression differentiates isolates showing early- and late-onset necrosis

We next aimed to quantitatively summarise the gene expression changes over time for the *K* most important genes identified by the random forest classifier (where the value of *K* must be chosen). To do this we used the Mahalanobis distance from the point of infection to each subsequent time point. Figure 1 shows the logarithm of Mahalanobis distances of the *K* = 10 most important genes at varying times. We see that these genes change significantly over the first two weeks (relative to the initial state) before plateauing or indeed decreasing. The consistency of this structure across isolates suggests that the Mahalanobis distance is a compelling measure to summarise gene expression changes during infection.

**Figure 1.**
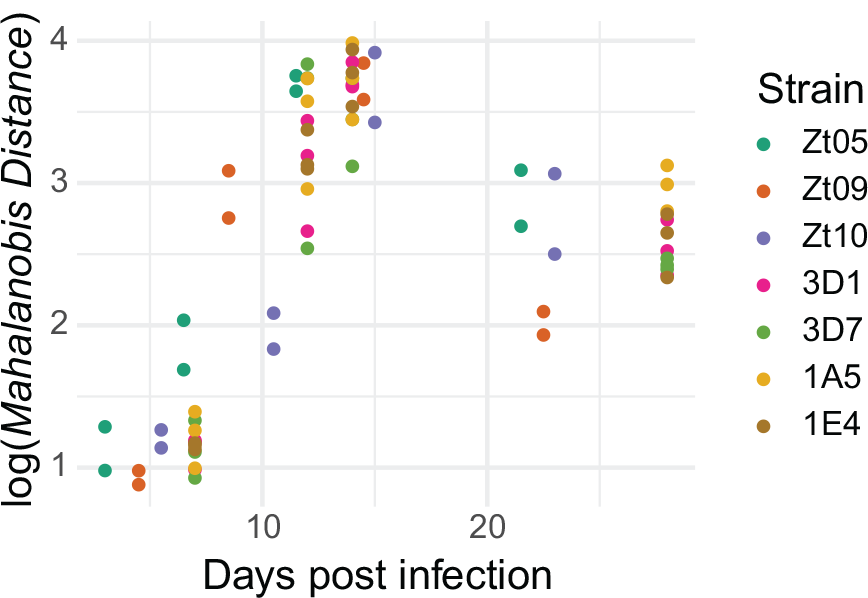
The log-Mahalanobis distance of gene expression, relative to the initial time point, for each time point, replicate and isolate. This analysis uses only the *K* = 10 most important genes identified by the random forest classifier that identifies those genes that change their expression significantly between the initial time point and the time point closest to necrosis development.

However, as seen in Figure 1, it is not immediately obvious which isolates progress more slowly or quickly, due to large gaps in the time series of observations, and because of misalignment between the timings of observations. Adopting the average timing of the ‘peak’ distance for each isolate as a proxy for the speed of infection, we can’t be certain if a peak coincides exactly with the timing of observations or instead lies within any of the gaps. We therefore adopt a Bayesian statistical model which makes simple assumptions about the functional form of the distance trajectory, to reduce the problem to a few parameters for each strain. Specifically, the model assumes the log-distances for each isolate can be described by a piece-wise linear trend plus an error term that captures variability between replicates. One of the parameters for each strain is the timing of the peak distance, which can be directly estimated along with measures of uncertainty.

Before proceeding, we must choose a value of *K* which determines how many important genes we use to compute the Mahalanobis distances. Here, the value of *K* can be thought of as a parameter, one which will inevitably affect the predictive performance of the statistical model: if *K* is too low, then we may be excluding genes which are informative for differentiating between early and late onset isolates; if *K* is too high, then we risk diluting the signal from the most informative genes. In this case, we believe the data available to us are insufficient to make a serious attempt to choose the optimal value of *K* without risking over fitting. We therefore discuss results in the context of different values of *K*: 5, 10, 15, and 20.

Figure 2 shows the posterior median expected log-distances and Figure S1 shows the posterior distributions of the timing of the peak distances, for each isolate and for different values of *K*. Across these values of *K*, there are some consistent patterns. First, the isolate with the longest time until necrosis development, Zt10, consistently has the latest peak distance. The other late onset isolate, 3D1, is also among the isolates with the latest peak distance. On the opposite end of the spectrum, the isolates with the earliest necrosis development, Zt05 and Zt09, are consistently among the isolates with the earliest peak distances. Although these distributions show some uncertainty, the consistent differences in early- and late-onset infection isolates suggest that our results are somewhat robust to the value of *K*. However, for these and later results it should be considered that, if the timings of the peak distances of the 7 isolates were ordered completely at random, there would be a just-below 5% probability that the two “late-onset” isolates would have the two latest peak distances (by chance). There is therefore a small but non-trivial chance that our results are random fortune. We fully acknowledge that to properly assess the significance of these results we would need a larger number of time points and isolates. Other substantive limitations of our analysis are discussed in the Methods.

**Figure 2.**
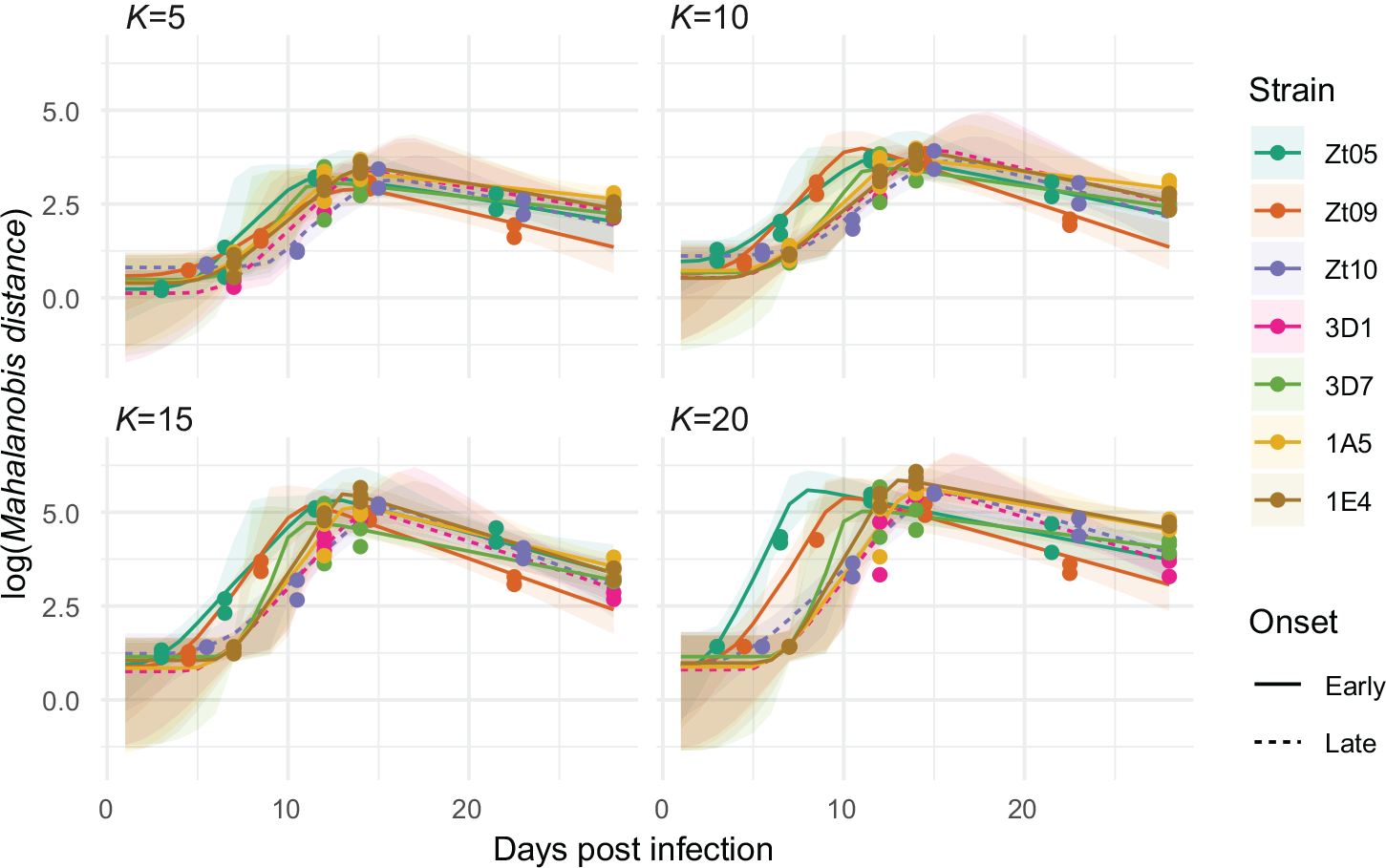
Posterior median expected log-Mahalanobis distance with 95% uncertainty intervals, for each isolate. Dashed lines emphasise the two late onset isolates Zt10 (purple) and 3D1 (pink), whose peaks appear delayed compared to the early onset isolates.

### Distinguishing early- and late-onset of infection for ‘novel’ isolates

Next, we used leave-one-out experiments to investigate the generality of our approach when applied to novel isolates. First, we investigated the stability of the important gene ranks across 7 runs of the procedure, where each one of the isolates was left out of gene selection in turn. Table 2 shows the 20 most important genes in order of median rank across the 7 experiments, as well as the lowest, mean and highest rank each gene achieved. Notably, one gene, Zt09_chr_1_00237 (My-cgr3G78133), was the most important in all 7 experiments. Many of these genes are also present in Table 1, with three exceptions; Zt09_chr_5_00763 (My-cgr3G72602), Zt09_chr_2_01239 (Mycgr3G69186) and Zt09_chr_13_00010. Fitting the parametric model once when each isolate is treated as novel, results in 7 distinct model fits. To summarise across these different fits, Figure 3 shows the expected log-Mahalanobis distance over time for each isolate, where the fit for each isolate is from the run where that isolate was treated as novel (using the *K* = 10 most important genes in each case). Here, we are looking to see if the patterns seen with respect to the timing of the peak distance hold, as shown in Figure 2. Indeed, we find that the peak distance for the two late onset isolates (Zt10 and 3D1) appear later in time than the early onset isolates, indicating that we can potentially distinguish between early- and late-onset of necrosis when we treat each isolate as unseen in turn.

**Table 2.** The 20 most important genes according to median rank across the 7 leave-one-out experiments, as well as their lowest, mean (rounded) and highest ranks.

| Median Rank | Gene ID | Reference ID | Gene Name | Lowest Rank | Mean Rank | Highest Rank |
| --- | --- | --- | --- | --- | --- | --- |
| 1 | Zt09_chr_1_00237 | Mycgr3G78133 | - | 1 | 1 | 1 |
| 2 | Zt09_chr_3_00489 | Mycgr3G91684 | - | 2 | 2 | 3 |
| 5 | Zt09_chr_9_00376 | - | - | 3 | 8 | 18 |
| 8 | Zt09_chr_5_00416 | Mycgr3G109517 | - | 4 | 10 | 23 |
| 9 | Zt09_chr_8_00269 | Mycgr3G105409 | - | 2 | 16 | 52 |
| 9 | Zt09_chr_2_00249 | Mycgr3G103091 | - | 3 | 14 | 40 |
| 10 | Zt09_chr_11_00021 | Mycgr3G49733 | - | 4 | 9 | 15 |
| 10 | Zt09_chr_12_00206 | Mycgr3G111474 | - | 4 | 13 | 32 |
| 13 | Zt09_chr_3_00873 | Mycgr3G70294 | - | 7 | 18 | 43 |
| 14 | Zt09_chr_1_00078 | Mycgr3G106752 | - | 7 | 19 | 40 |
| 15 | Zt09_chr_1_00619 | Mycgr3G65617 | - | 6 | 14 | 24 |
| 16 | Zt09_chr_3_00461 | Mycgr3G103692 | - | 2 | 14 | 22 |
| 16 | Zt09_chr_1_00142 | Mycgr3G106779 | MgEXG1 | 4 | 17 | 37 |
| 18 | Zt09_chr_12_00203 | Mycgr3G50464 | MgAtr2 | 7 | 22 | 46 |
| 25 | Zt09_chr_3_00393 | Mycgr3G99503 | - | 8 | 25 | 39 |
| 26 | Zt09_chr_13_00010 | - | - | 21 | 32 | 49 |
| 27 | Zt09_chr_2_01239 | Mycgr3G69186 | - | 5 | 27 | 54 |
| 28 | Zt09_chr_6_00030 | Mycgr3G109703 | - | 4 | 30 | 50 |
| 30 | Zt09_chr_5_00763 | Mycgr3G72602 | - | 15 | 36 | 55 |
| 30 | Zt09_chr_7_00076 | Mycgr3G73671 | - | 2 | 31 | 75 |

**Figure 3.**
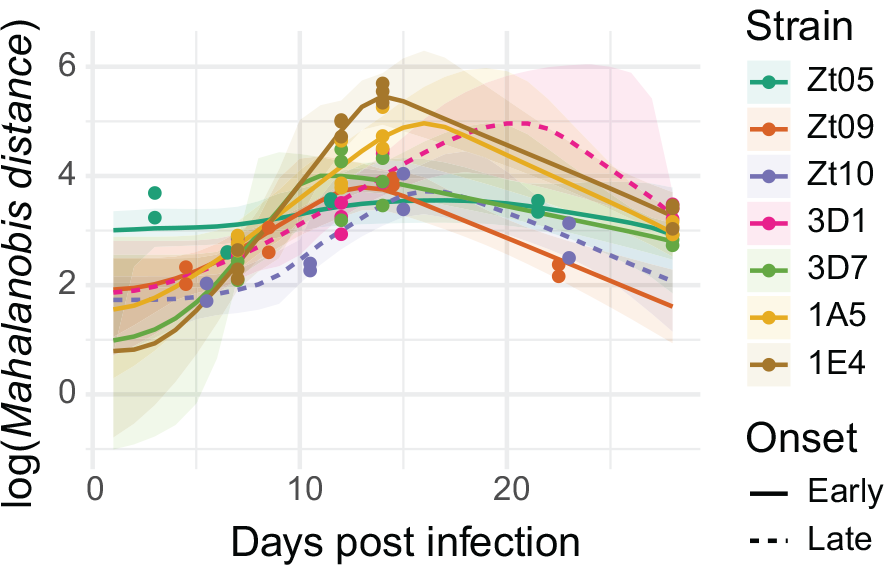
Posterior median expected log-Mahalanobis distance with 95% uncertainty intervals, where the fit for each isolate is from the selection and model run where that isolate was treated as novel with all four time points and using the *K* = 10 most important genes. Different colours represent the isolates and the dashed lines emphasise the two late onset isolates Zt10 (purple) and 3D1 (pink). Noticeably the peaks in posterior median expected log-Mahalanobis distance are delayed in the late onset isolates.

### Predicting the timing of infection milestones for ‘novel’ isolates with limited biological measurements

Finally, we also investigated whether our approach could be used to predict the timing of the peak Mahalanobis distance of a novel isolate, and by extension, the timing of infection milestones without observing the whole time series for that isolate. To do this we adjusted our leave-one-out experiment so that the distances at the third and fourth time points for the novel isolate were treated as missing values. This means that the estimation of the timing of the peak distance (from the Bayesian statistical model) is only based on the first two time points for the novel isolate. This would demonstrate the potential of our methods for predicting the virulence of new pathogenic isolates with few biological measurements.

First, we assessed how well our model performs when extrapolating beyond the available time points by comparing the distance values we treated as missing to their associated predicted values. Figure 4 shows posterior median predictions and 95% prediction intervals for the third and fourth time points. From this plot we can see that predictions are more accurate for the 3rd time point – which is arguably more important than the 4th, as it is closest to the peak we are trying to predict. Indeed, the majority of variability in the true novel log distances is explained by the predictions (*R*^2^ = 0.71) for the 3rd time point. We can also see that the 95% prediction intervals cover the observed values almost every time. There is some evidence that the model may systematically under-predict the distance at the 3rd time point, which should be taken into account when designing future versions of this approach.

**Figure 4.**
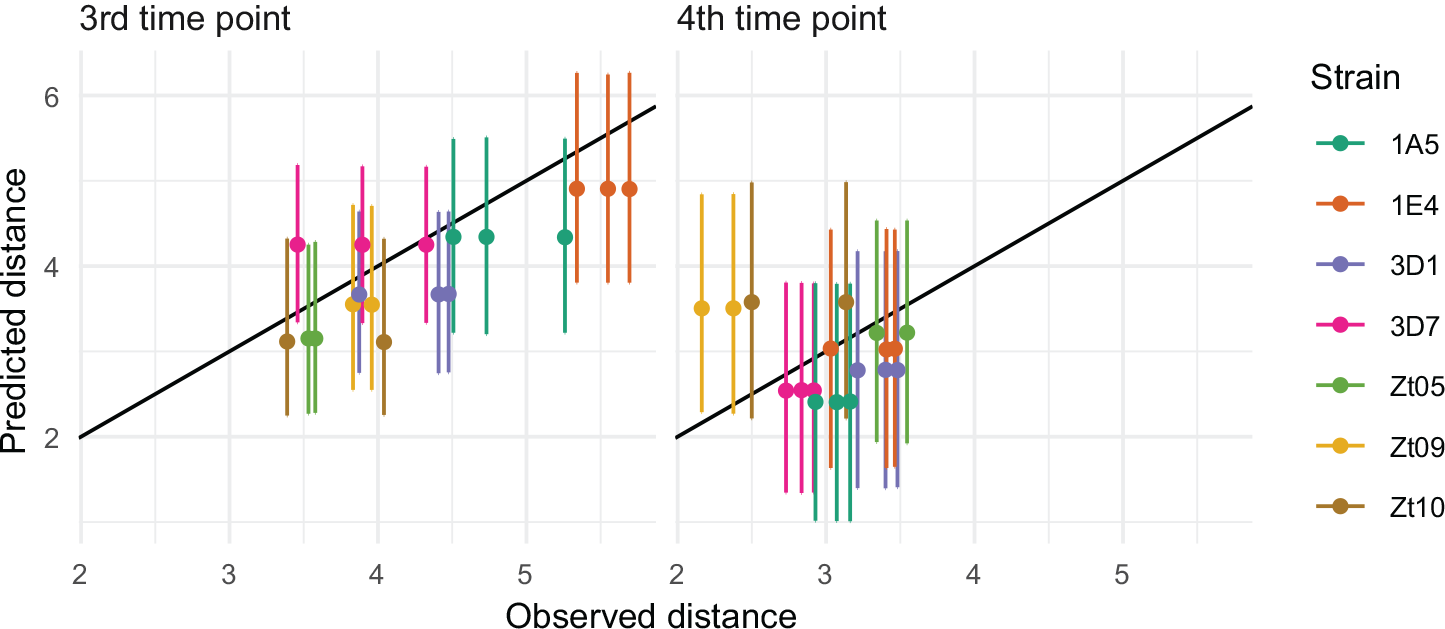
Posterior median predictions of the log-Mahalanobis distance at the 3rd and 4th time points of the novel isolates (which were treated as missing values), with 95% uncertainty intervals, using the *K* = 10 most important genes. Different colours represent the isolates.

Figure 5 shows the posterior median expected log-Mahalanobis distance where the fit for each isolate is taken from the model where that isolate was treated as novel (and therefore the 3rd and 4th time points were treated as missing values). Here the peak distance for the late onset isolates is noticeably later than for the early onset isolates, implying that when using this approach there is potential to predict the onset of infection for a new isolate when using only existing isolates for feature selection and when only observing the first portion of the time series for the new isolate. These results demonstrate the potential utility of our approach for predicting infection outcomes, in this case the onset of necrosis, for new isolates using only limited biological measurements. In turn, these results therefore have implications for the use of these methods for developing new diagnostics and forecasting the emergence of new disease.

**Figure 5.**
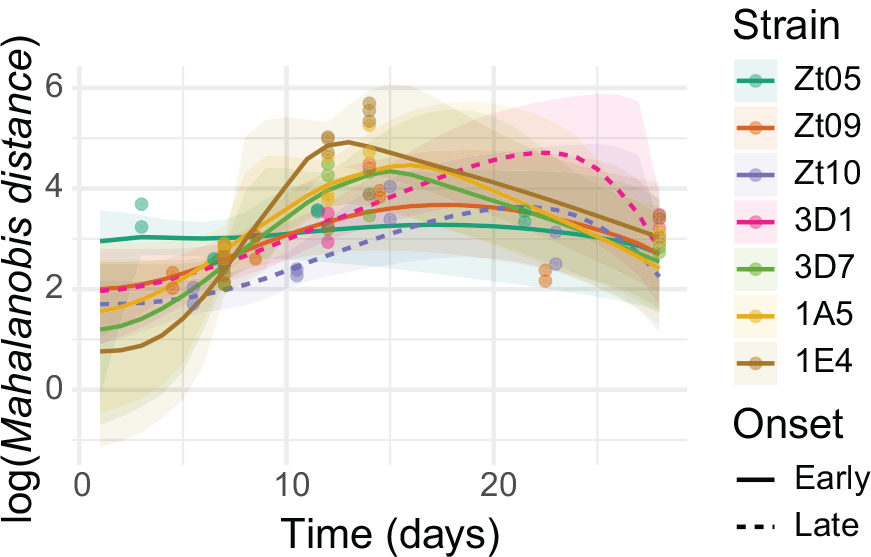
Posterior median expected log-Mahalanobis distance with 95% uncertainty intervals, where the fit for each isolate is from the selection and model run where that isolate was treated as novel using only the first two time points and the *K* = 10 most important genes. Different colours represent the isolates and the dashed lines emphasise the two late onset isolates Zt10 (purple) and 3D1 (pink). Noticeably the peaks in posterior median expected log-Mahalanobis distance are delayed in the late onset isolates.

### Bioinformatic analysis of selected most important genes

Tables 1 and 2 report the most important genes predictive of infection timing. Using the Gene Ontology (GO) [89] it is possible to identify three common functions for these genes (i) transporters, (ii) transcriptional regulators and (iii) enzymes, particularly oxidoreductases. Zt09_chr_12_00203 (MgAtr2) is an experimentally validated ATP binding cassette (ABC) transporter [90], Zt09_chr_1_00142 (MgEXG1) is a putative exo-beta-1,3-glucanase and Zt09_chr_- 6_00030 (My-cgr3G109703) is described as a putative major facilitator superfamily transporter [25]. Other potential transporters can be identified using GO annotation; Zt09_chr_6_00030 (Mycgr3G109703), Zt09_chr_4_0050 (Mycgr3G39898) and Zt09_chr_12_- 00206 (Mycgr3G111474) are all annotated with the transmembrane transport GO term (GO:0055085). Zt09_chr_1_00142 (MgEXG1), Zt09_chr_10_00159 (Mycgr3G105887) and (Mycgr3G106752) are associated with transcriptional regulation, and Zt09_- chr_7_00076 (Mycgr3G73671) is annotated with the DNA-binding transcription factor activity (GO:0000981) GO term. Many genes have enzymatic functions, particularly oxidoreductase activity. Zt09_- chr_12_00206 (Mycgr3G111474) is described as being involved with the sphingolipid metabolic process (GO:0006665) and as having oxidoreductase activity (GO:0016491). Zt09_chr_3_00393 (Mycgr3G99503) is annotated as having oxidoreductase activity, acting on NAD(P)H (GO:0016491). Zt09_chr_8_00269 (Mycgr3G105409) is annotated with response to oxidative stress (GO:0006979) and oxidoreductase activity (GO:0016491). Zt09_chr_3_00873 (Mycgr3G70294) is inferred to be involved in the oxidative-reduction process (GO:0055114) and as having choline dehydrogenase activity (GO:0008812). Zt09_chr_5_- 00763 is annotated as being involved with lipid biosynthesis (GO:0008610) and oxidation-reduction processes (GO:0055114), as well as having oxidoreductase activity (GO:0016491). Finally, no annotation information is available for the remaining genes Zt09_chr_1_00237 (Mycgr3G78133), Zt09_- chr_2_00249 (Mycgr3G103091), Zt09_chr_3_00489 (Mycgr3G91684), Zt09_chr_8_0027 (Mycgr3G110409), Zt09_chr_2_01239, Zt09_chr_9_00376 and Zt09_chr_- 13_00010 with the latter 2 genes not appearing in the reference genome [25], indicating they could be specific to the isolates analysed in this study.

In order to expand on the qualitative functional descriptions of the most important genes, we next performed enrichment analysis for these genes in *Z. tritici* and of orthologs in other well annotated fungal species. In *Z. tritici*, we found enrichment for GO terms and pathways relating to lipid and sphingolipid biosynthesis processes, sphingolipid desaturase activity and oxidoreductase activity. However, although these enrichments were significant (*p-value <* 0.05), none were significant after false discovery rate correction (Table S2). The first set of orthologs analysed were in another Dothideomycete, *Cladosporium fulvum*. We used the MycoCosm database [91] to identify orthologs and to find GO, InterPro and EuKaryotic Orthologous Groups (KOG) annotations. Although the MycoCosm database doesn’t allow the identification of enrichment of annotations, we find that the annotations match closely what was observed in *Z. tritici*. GO terms relating to oxidoreductase activity (GO:0016651), ATP binding/ATPase activity (GO:0005524, GO:0016887) and transporter activity (GO:0005215, GO:0042626) are all associated with *C. fulvum* orthologs. These patterns are repeated for InterPro and KOG annotations where oxidoreductase subunits (IPR011538, IPR011537, IPR001949) and ABC transporter-like domains (IPR003439, IPR013525, IPR010929) are well represented. KOG annotations show similar patterns with transcription factors (KOG2294), oxidoreductase subunits (KOG2658) and ABC transporters, related to drug resistance (KOG0065), all present (Table S3). Next, we chose another wheat pathogen, *Fusarium graminacearum*, and identified orthologs and enriched GO terms and pathways using FungiDB [92]. As before, we find annotations for a variety of GO terms related to transport (GO:0015168, GO:0015105, GO:0042626, GO:0015399, GO:0022804, GO:0015103), oxidoreductase activity (GO:0016655, GO:0016491, GO:0055114, GO:0072593) and sphingolipid biosynthesis (GO:0042284, GO:0030148) are enriched but not significant after false discovery rate correction. However, we find several Cellular Component GO terms are significantly enriched (*corrected p <* 0.05) relating to the cell membrane (GO:0005743, GO:0019866, GO:0005886, GO:0031966, GO:0005740, GO:0016020, GO:0031975, GO:0031967), supporting the suggestion that these genes may be functionally related to transport across the cell membrane (Table S4). Finally, we analysed orthologs and their functions in the well-studied, model filamentous fungi, *Aspergillus fumigatus* (Table S5). To support previous findings, GO terms relating to oxidoreductase activity (GO:0016491) and cell wall organisation (GO:0070871, GO:1904541, GO:0044277, GO:0071853) are significantly enriched (*corrected p <* 0.05). Functional pathway analysis of the most important *Z. tritici* genes and their orthologs supports the observation that these genes are likely to be involved in enzymatic reactions, particularly oxidation-reduction reactions, and transport, potentially across the cell membrane.

## Discussion

Machine learning methods have recently been applied to the study of plant-pathogen interactions in the areas of disease monitoring, genomic selection for resistance and prediction of potential effectors (reviewed in [82]). In this study we apply a multi-stage machine learning approach with a parametric statistical model to publicly available transcriptomic data for several isolates of the wheat pathogen *Z. tritici*. Our methods are capable of identifying a subset of genes that significantly change their expression between the onset of infection and an infection outcome measured as the first appearance of necrosis. These genes include those with disease-related functions and as yet uncharacterised or isolate-specific genes that might represent novel genes with roles in disease. We next find that the changes in expression of these genes, summarised by the Mahalanobis distance, differentiates between fungal isolates that show an early or late appearance of necrosis. Finally, we demonstrate potential ability to discriminate between “early” and “late” necrosis onset of novel isolates.

It has been proposed that machine learning methods will be central to analysing high-resolution omics data, such as those from multiple members of a species to understand plant-pathogen interactions [93]. However, current applications of machine learning methods to transcriptomics data of plant-pathogen interactions focus largely on predicting stress conditions [83], genes associated with specific traits [84] or predicting gene regulatory networks (GRN) in the host [94]. For fungal pathogens, transcriptomic datasets have been used to infer a GRN for *Fusarium graminearum*, with the aim of identifying major regulators of disease pathways that provide candidates for experimental validation [95]. Our work represents, to the best of our knowledge, the first approach that aims to use high-resolution transcriptomics data from multiple members of a species to predict infection outcomes and, in doing so, identify those genes most predictive of that infection outcome, in this case the onset of necrosis.

Identifying the most important genes that predict the onset of necrosis may yield useful insights into the disease process and provide novel candidates for forecasting the virulence of new isolates. In general, those genes found to be important in predicting disease fall into three categories; (i) transporters, (ii) transcriptional regulators and (ii) enzymes, particularly oxidoreductases (Tables 1 & 2). These functions are supported by pathway analysis of the *Z. tritici* genes (Table S2) and orthologs in other relevant fungal species (Tables S3-S5). Zt09_- chr_12_00203 (MgAtr2) is a known ABC transporter that provides protection against toxic compounds including fungicides and plant metabolites [90, 96], but is not essential for virulence [97]. Zt09_chr_6_00030 (My-cgr3G109703) is a putative major facilitator superfamily transporter and other members of this family, such as MgMfs1, have been shown to confer protection against natural toxic compounds and fungicides in *Z. tritici* [98]. In addition to transporters, many of the important genes are annotated as having oxidoreductase activity and are potentially involved in the response to oxidative stress. Responding to reactive oxygen species, which plays a role in host defense, is important to both fungal pathogens of plants [99] and humans [100]. For example, Zt09_chr_12_00206 (Mycgr3G111474) is annotated as being involved in the sphingolipid metabolic process (GO:0006665), where it has been speculated that microbes containing sphingomyelin would be more resistant to damage by oxidative stress [101], with implications for fungal pathogenesis [102, 103]. Many of the genes identified as being most important for the transition to necrosis are uncharacterised and, as such, represent new predictions of genes with roles in disease and potential biomarkers for disease forecasting and surveillance. Previous work applying machine learning techniques to infectious disease have identified the most important mutations that predict the human transmission of avian influenza viruses with implications for disease monitoring [104]. To differentiate between isolates showing early and late onset of necrosis development we summarised the changes in expression of the most important genes using the Mahalanobis distance. A Bayesian parametric model applied to the distances then correctly distinguishes early and late onset for varying numbers of most important genes (*K*) (Figure 2). Furthermore, we have shown that this measure has potential promise in predicting infection timings of ‘novel’ isolates using leave-one-out experiments (Figure 3) and with only limited timepoints from the transcriptomics series (Figures 4 & 5). We demonstrate the utility of our approaches to predicting the onset of necrosis development in new isolates of *Z*.*tritici* using only the most important genes identified in this study and limited data on infection collected for the new isolate. The results indicate that the methods presented here could be used to predict the impact of a new pathogenic isolate. Previous work in related fields has focused on assessing pre-planting factors, such as latitude and longitude, to predict the susceptibility of wheat to outbreaks of *Parastagonospora nodorum* [105]. Studies have also used machine learning on images, fluorescence and thermography to classify leaves as infected or uninfected [106] and predict disease development stages [107]. All these studies use machine learning methods for the prediction of disease, however our approach is the first to use pathogen transcriptomics as the predictor variables rather than imaging. This study adds further evidence for the potential applicability of machine learning approaches for the prediction of disease outbreaks that may inform disease management decisions.

Although we have potentially succeeded in deploying machine learning approaches to predict the onset of necrosis in an important fungal pathogen, our approach is somewhat limited by data availability and inconsistency. The publicly available RNA-Seq data used in this study has been collected based on different criteria, with one study collecting data at pre-defined time-points [31] and another at different disease development stages [30]. Consistent transcriptomic data collection, as well as more fine-grained expression data, would further increase the predictive accuracy of the proposed approach. Likewise, the infection outcomes available for this study also limited our predictions to the onset of necrosis. This highlights the need for quantitative information on disease progression, which is becoming increasingly more feasible with the development of low-cost phenotyping imaging setups and software for automated image analysis [108], though additional work is still required before these approaches can be used to quantitatively measure fungal infection.

The approach outlined in this study takes a noteworthy step towards demonstrating the promise of machine learning to identify the most important genes predictive of infection outcomes. In this application, many of the most important genes are uncharacterised and thus, this approach may be useful for the annotation of genes with roles in disease and to identify biomarkers that can be used to develop new diagnostics for disease monitoring and forecasting. Moreover, we have demonstrated that we are capable of predicting a disease outcome for novel isolates using only the first two gene expression timepoints, which has further implications for the speed and ease of data collection to deploy these approaches for disease surveillance and prediction. We can envisage a scenario where these approaches can be applied to a newly emerging isolate of a known pathogenic species, where existing data could be used to train our models and the collection of minimal new data (we have demonstrated our approach on transcriptomics data of only 2 infection timepoints) on the emerging isolate can be used to predict its infectivity. Of course, there are many potential avenues for future work including the integration of different omics data [82], reduction of the high-dimensionality of gene expression data with network biology approaches [109] and use of automated imaging data to capture infection phenotypes [108], that may reduce data collection and broaden the phenotypes that can be predicted. Together, these advancements are a significant step towards wider utility and application of machine learning methods to study plant-pathogen interactions with implications for developing new monitoring techniques, control strategies and treatments.

## Methods

### Data gathering and pre-processing

We used gene expression data from two studies; *(i)* Haueisen *et al*. characterised gene expression for 3 isolates of *Z. tritici* (Zt05, Zt09 and Zt10) at 4 stages of infection (A, B, C and D) determined by confocal microscopy on wheat cultivar Obelisk, each with 2 replicates [30]. Infection stages can be converted into dpi for each isolate using the results of the confocal microscopy experiments reported by Haueisen *et al*.. Note, Haueisen *et al*. reported ranges for the timing of each infection stage, and we took the mid-point of these ranges. These data were downloaded as raw counts from the Gene Expression Omnibus [110] under accession GSE106136. *(ii)* Palma-Guerrero *et al*. recorded gene expression data for 4 isolates of *Z. tritici* (3D7, 3D1, 1A5 and 1E4) infecting wheat cultivar Drifter at 7, 12, 14 and 28 dpi with 3 replicates [31]. These data were downloaded as raw counts provided in Supplementary Table 2 of the original publication. Importantly, sequencing data from both datasets were aligned to the Zt09 annotation [78] meaning all isolates were annotated with consistent gene identifiers.

To normalise raw count data to account for gene length and library size (number of reads mapped to genes) we used Reads Per Kilobase of transcript per Million mapped reads (RPKM). Gene lengths for the Zt09 annotation were taken from Haueisen *et al*. [30] and library size was defined as the total number of reads assigned to genes in each sample. In each dataset, genes were removed if there were no expression values for a gene in all replicates in any single timepoint. For any timepoint with only partial data, the missing data was filled using the mean of the expression values of the other replicates for that isolate and timepoint. Filtering left 9,371 and 6,641 genes in the Haueisen *et al*. and Palam-Guerrero *et al*. data respectively. Normalised data were reported as log(RPKM + 1) and are available in Supplemental Tables S6 and S7 for the data from Haueisen *et al*. and Palma-Guerrero *et al*. respectively.

### Infection outcomes

Both Haueisen *et al*. and Palma-Guerrero *et al*. report a variety of infection outcomes including the timing of disease symptom development such as necrosis and pycnidia and the percentage leaf area covered by necrosis (or necrosis level) and pycnidia. For this study we used the most comparable measure common to both experiments, which was the timing of the appearance of necrosis measured in dpi. We used the timing of the appearance of necrosis to broadly classify infection by these isolates as early or late onset. This follows the findings that isolate Zt10 develops necrosis significantly later than isolates Zt05 and Zt09 [30]. Symptoms of infection, including necrosis were also found to occur later and be less severe for isolate 3D1 compared to 3D7, 1A5 and 1E4 [31]. Supplemental Table S8 shows the infection timing and classification for each isolate used in this study.

### Differential expression analysis

In order to be able to compare the results of our machine learning approach with the more traditionally used differential expression analysis, we identified significantly differentially expressed genes using edgeR [88]. Briefly, we merged the Haueisen *et al*. and Palma-Guerrero *et al*. data sets on common gene IDs and used edgeR’s default methods for filtering and normalisation of count data. We defined groups to compare the first infection timepoint (defined as infection stage A and 7 dpi in the Haueisen *et al*. and Palam-Guerrero *et al*. data respectively) to the timepoint of ‘peak’ Mahalanobis distance (defined as infection stage C and 14 dpi in the Haueisen *et al*. and Palam-Guerrero *et al*. data respectively - see below) in those isolates classified as early infecting. The exact test was used to identify significant differential expression with *p −values* corrected by the method of Benjamini and Hochberg [111]. Note that this set of differentially expressed genes is used only for comparison purposes, but it is not otherwise used in the proposed approach.

### Gene selection

The first stage of our approach is to identify a small subset of genes which change significantly over the infection period and therefore may be candidate genes important in the disease process. To do this we apply a machine learning (random forest [112]) classifier to differentiate between observations of gene expression made close to the infection time and observations made close to the time of necrosis development. Specifically, the ‘output’ for time *t*, replicate *r* and isolate *s*, denoted by *y*_*t,r,s*_, is defined as a binary quantity where *y*_*t,r,s*_ = 0 if *t* is the initial time point of the experiment and *y*_*t,r,s*_ = 1 if *t* is the time point closest to necrosis development. This time is taken to be stage C in the Haueisen experiment and 14 days in the Palma-Guerrero experiment. For each output *y*_*t,r,s*_ we also have a corresponding vector of gene expressions ***x***_*t,r,s*_ which serves as the input to the classifier. The main result of doing this is not the fitted classifier itself but the ability to quantitatively assess how predictive each gene is in determining at what stage of infection the observation is made. As such a ranking of genes by importance is obtained by computation of the impurity score for each gene and sorting the scores into descending order. The gene with the highest score is therefore the most ‘important’, and so on. It then remains to choose a value of *K* whereby we carry forward only the *K* most important genes into the subsequent stages.

Random forest classifiers are inherently stochastic, meaning the ordering of gene importance can vary when the model is fit several times to the same data. To improve the stability of the importance ranking, we used an ‘extra trees’ version of the random forest classifier (R package ranger [113]) and employed a large number of trees (100k).

### Mahalanobis distance

The second stage of our approach is to quantitatively summarise the change in expression over time of a subset of relevant genes. That is, we want to characterise, with a single numerical value, the state of the biological system with respect to the infection process, so that, ideally, a state of infection is clearly distinguishable from the non-infected state. To accomplish this, we adopt all observations of these genes made at the first time point in each experiment as a baseline. Using these observations, we compute a mean expression vector ***µ*** and an empirical covariance matrix **Σ**. Then, for observations made at time *t*, of isolate *s* and of replicate *r*, denoted by ***x***_*t,s,r*_, the distance *d*_*t,s,r*_ *>* 0 from the baseline distribution (as summarised by ***µ*** and **Σ**) is computed using the Mahalanobis distance. For given, ***x, µ*** and **Σ**, this distance is defined by^1^:

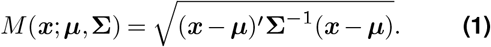

The notion of Mahalanobis distance allows us to take into account the natural variability of each gene via their estimated dispersion and to treat correlated genes in an appropriate way. Rather than treating each gene independently, the Mahalanobis distance allows us to distinguish when the changed expression level of a particular gene is contributing to characterise a diverse state or when, given that other correlated genes behave in a similar fashion, its contribution should be discounted. By limiting the information to a small subset of discriminative genes identified in stage one, we significantly improve the signal to noise ratio and avoid the confounding effects of genes that are not related to the infection process (e.g. housekeeping genes).

### Parametric model

We next aimed to investigate whether the timing of the ‘peak’ Mahalanobis distance might give an insight into the relative timings of certain infection milestones, such as the development of necrosis. There is no guarantee, however, that the timing of observations will directly coincide with the peak distance. Moreover, given observations at a limited number of fixed time points, it is not possible to pinpoint the timing of the peak precisely. Therefore, we seek to estimate the timing of the peak for each isolate, with associated uncertainty, in a way which generalises across experiments that make observations at different times post infection. To achieve this we adopt a relatively simple parametric statistical model in the Bayesian framework. This approach also offers the potential to predict the timing of a novel isolate’s peak (and therefore infer its infection characteristics) without observing the whole time series. For a given isolate *s* observed at time *t*, the model assumes independent, identically distributed Normal (Gaussian) errors for the logarithm of the Mahalanobis distance *d*_*t,s*_:

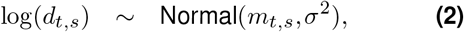

where *m*_*t,s*_ is a piece-wise linear function of time for each isolate and *Σ* captures variability associated with observation error and differences between replicates. The functions *m*_*t,s*_ are defined as linear in between four ‘knots’ which are different for each isolate. Each knot has a corresponding x-axis (time) value, and a y-axis (log-Mahalanobis distance) value. Figure S2 is a visual aid for understanding the structure of this function. For a particular isolate, the first knot in time is fixed at 0 days post infection and shares a log-distance value with the second knot. The second knots, seen as the first break-points following flat periods in Figure S2, therefore represent the time when the expression of the most important genes begin to increase significantly. We will therefore refer to the second knots as the ‘point of increase’. The third knots then represent the ‘peak distance’, and the final (fourth) knots, fixed at 28 days, represent the distance at the ‘end point’ of the experiment. For a particular isolate, we therefore have two knots (‘point of increase’ and ‘peak distance’) where the timings (x-axis coordinates) are largely free parameters. In the Bayesian framework for statistical inference, we must specify ‘prior’ distributions for all parameters, representing our uncertainty about those parameters in the absence of any data. To be as uninformative as possible, we define the timing of these knots as parameters with uniform prior distributions spanning 0 to 28 days, with only the natural constraint that the peak must occur after the point of increase. We then have three knots (‘point of increase’, ‘peak distance’ and ‘end point’) where the log-distances (y-axis coordinates) are largely free parameters, which we denote by ***α*** = *α*_1,*s*_, *α*_2,*s*_, *α*_3,*s*_ in order of increasing time. Here we specify that the log-distance (y-axis value) of these knots for different isolates come from a common distribution. This is achieved by treating the *α*_*j,s*_ as random effects:

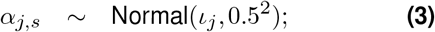

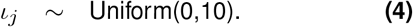

For the point of increase, peak distance and the end point knots, *ι*_1_, *ι*_2_, *ι*_3_ define the expected log-distance, respectively. In summary, the timing of the knots is independent across isolates, but the distance is similar. Using this specification, it is possible to fit observations made at arbitrary time points and make predictions of the distance for a particular isolate both in-between and beyond the observed time points. In the Bayesian framework, the observations update the prior distributions, resulting in posterior distributions which quantify our uncertainty in the model parameters. We employ the Markov chain Monte Carlo (MCMC) method (R package nimble [114]) to obtain samples from the joint posterior distribution of all parameters, subject to convergence criteria being met. For any given parameter, we can derive central point estimates (e.g. by computing the mean of the posterior samples) and measures of uncertainty (e.g. 95% intervals can be obtained by computing the 2.5% and 97.5% quantiles of the samples). Using the Monte Carlo simulation method, we can then obtain samples from the posterior predictive distributions for missing or new data points. These predictions jointly account for both random variability as quantified by the model and uncertainty in the model parameters.

Figure S2 shows the posterior median expected log-Mahalanobis distance defined by the piecewise linear functions for each isolate, with 95% uncertainty intervals. Dashed lines are used to distinguish the two ‘late’ onset isolates, Zt10 and 3D1. Notably, although the model is specified as piecewise linear given the positions of the knots, treating these positions as random variables smooths the functions considerably once parametric uncertainty is taken into account. Looking at the estimated functions, the peaks for the two late onset isolates appear to occur later than all the ‘early’ onset isolates. This can be investigated more clearly by analysing the posterior distributions for the timing of the peak knots, shown in Figure S3. As the number of replicates and time points for each isolate is quite small, these distributions are quite uncertain. Moreover, they appear quite misshapen compared to most analysed posterior distributions, despite MCMC convergence and the number of posterior samples being relatively high. If more data were available, a potential and indeed straight-forward extension to the model would be to directly relate the parameters controlling the timing of the peaks with covariates such as the timing of necrosis, the timing of pycnidia appearance, or the necrosis level. For seen isolates the values of these covariates would be known, but would be left as missing values for novel isolates. Using the Bayesian approach, these infection outcomes could then be predicted - assuming the relationships between the timing of the peak and these covariates turn out to be reasonably strong - to allow for a richer assessment of a novel isolate’s infection capability.

### Model validation

Up to this point we have explained our potential for identifying a small subset of important genes and, to uncover relationships between the expression of these genes and infection timings. However, we have so far treated all of the available data as known. To investigate whether we can also predict how quickly a novel isolate will reach infection milestones, we have employed a leave-one-out approach where each of the 7 available isolates is treated as novel in turn. For each isolate the process is as follows:

Step 1: Perform gene selection with the random forest classifier using only the 6 seen isolates as data.

Step 2: Calculate Mahalanobis distances for all 7 isolates but using only the 6 seen isolates as inputs for calculating ***µ*** and **Σ**.

Step 3: Fit the parametric model to all 7 isolates twice, first with data for all isolates and all time points (a) and second using only the first two time points for the novel isolate (b).

Withholding the novel isolate from gene selection ensures that any conclusions drawn from the timing of the peak distance relies only on the important genes from the other 6 isolates. Meanwhile, by fitting the parametric model a second time with only the first two time points for each novel isolate, we can investigate the potential of our approach for predicting the timing of the peak distance, and by extension how quickly the infection will develop, without observing the whole time series. In the latter case (b), we treat the distances for the novel isolate at the third and fourth time points as missing values. For these values samples can then be obtained from their respective posterior predictive distributions, available from the fitted Bayesian model.

In assessing these predictions, we are primarily interested in: i) the average error of predictions (bias); ii) the average absolute error of predictions (accuracy) and;

iii) the reliability of prediction intervals in capturing the true novel values (coverage). For i), when predicting the log() distances at the third time point, the average of all observed values minus all predicted values (posterior means) was 0.20, indicating slight under-prediction overall. When predicting the fourth time point, the average error was -0.08, indicating little bias relative to the scale of the input data. For ii), the average absolute difference between the observed values and predicted values was 0.5 for the third time point and 0.54 for the fourth. This is only moderately larger than the expected absolute error of 0.38 from the model with *K* = 10 and all isolates and time points observed (Figure S2), as quantified by the posterior mean of *σ*. When comparing predictions for the third time point to the true novel value, a substantial *R*^2^ = 71% of the variability in the true values is explained by the predictions. For the fourth time point, *R*^2^ is negative, indicating that the model can not effectively predict the Mahalanobis distance of novel isolates at the end point of the experiment, relative to other isolates. For iii), the 95% prediction intervals contained the true novel values approximately 95% of the time for both time points. Taken together, these metrics suggest that predictive point estimates for novel isolates and time points are reasonably accurate (relative to the expected error in the model with all isolates and time points observed) and that 95% prediction intervals are likely to contain the unknown true log distances.

### Bioinformatic analysis of selected genes

To further analyse the genes that change expression significantly and can be used to distinguish between early- and late-onset of infection, we performed pathway analysis and examined orthologs from other species. Pathway analysis was performed using FungiDB [92]. Briefly, we used FungiDB’s BLASTn search functionality to search the sequences of the most important genes against the *Z. tritici* IPO323 sequence to convert between IDs used in this study and those used by FungiDB. Next, we used the FungiDB search strategies to perform pathway analysis on these IDs to identify enriched GO terms, KEGG annotations and MetaCyc pathways. Annotations and pathways were deemed to be significantly enriched with a Benjamini and Hochberg [111] corrected *p-value < 0*.*05*.

In order to examine orthologs we chose 3 species, another Dothideomycete species *Cladosporium fulvum, Fusarium graminacearum* a wheat pathogen from the family Nectriaceae and *Aspergillus fumigatus* a well-studied model filamentous fungi. For *C. fulvum*, we used the MycoCosm database [91] BLAST functionality to identify *C. fulvum* orthologs to the selected *Z. tritici* genes. We next assembled the GO, InterPro and Eukaryotic Orthologous Groups (KOG) for the identified orthologs. We used FungiDB [92] to identify orthologs and conduct pathway enrichment analysis for *F. gram-inacearum* and *A. fumigatus*. As before, we used the FungiDB search strategies to perform pathway analysis on *F. graminacearum* and *A. fumigatus* orthologs to identify enriched GO terms, KEGG annotations and MetaCyc pathways. Annotations and pathways were deemed to be significantly enriched with a Benjamini and Hochberg [111] corrected *p-value < 0*.*05*.

### Substantive limitations of the results for distinguishing early versus late infection onset isolates

As mentioned in the results section, we acknowledge that the correct ordering of the peak distances, such that the two “late-onset” isolates have later peaks than the 5 “early-onset” isolates, could have arisen from random ordering, with a probability just under 5%. Other substantive limitations include:

- For the Haueisen *et al*. data, we derived dpi values (days post infection) for the infection stages by taking the mid points of the ranges they reported. We did not explore the sensitivity of our results to this choice.
- We applied our methods to genetic expression data subject to a log(RPKM + 1) transformation. We did not explore the sensitivity of our results to alternative transformations.

## Supporting information

Supplementary-file1-Diff-exp

Supplementary-file2-Cladosporium-fulvum

Supplementary-file3-Fusarium-graminacearum

Supplementary-file4-Aspergillus-fumigatus

Supplementary-file5-Haueisen

Supplementary-file6-Palma

Supplementary-file7-Onset

Supplementary-file8-pathway-analysis

## Data availability

Data are available both in supplementary data and via a GitHub repository, which includes the code used for this work (https://rmames.github.io/ml_zymoseptoria/ML_-Zymo.nb.html). An R notebook of this project is available via GitHub pages (https://rmames.github.io/ml_zy-moseptoria/ML_Zymo.nb.html).

## Author Contributions

OS was responsible for experimental design, data analysis and contributed to writing the manuscript. GT contributed to writing the manuscript. FC contributed to project conception, experimental design and writing the manuscript. RMA contributed to project conception, experimental design, data gathering and writing the manuscript.

## Funding

RMA and GT were supported by a BBSRC/EPSRC Interface Innovation Fellowship (EP/S001352/1). The funders had no role in study design, data collection and interpretation, or the decision to submit the work for publication.

## Conflict of interest

The authors declare that the research was conducted in the absence of any commercial or financial relationships that could be construed as a potential conflict of interest.

## Supplementary Information

**TABLE S1**. Supplemental file 1 contains lists of differentially expressed genes identified between the first infection timepoint and the timepoint of peak Mahalanobis distance. The columns include gene ID, log-fold change, log counts per million (CPM), *p − value* and false discovery rate corrected *p − value*.

**TABLE S2**. Supplemental file 2 contains the MycoCosm analysis results for *C. fulvum* orthologs of the most-important *Z. tritici* genes. The sheets include (i) BLAST results converting gene identifiers used in this study to *C. fulvum*, (ii) GO annotations of these genes, (iii) InterPro annotations of these genes and (iv) KOG annotations of these genes.

**TABLE S3**. Supplemental file 3 contains the FungiDB enrichment results for the analysis of *F. graminacearum* orthologs of the most-important *Z. tritici* genes. The sheets include (i) BLAST results converting gene identifiers used in this study to *F. graminacearum* identifiers, (ii) the results of GO enrichment analysis for the MF ontology, (iii) the results of GO enrichment analysis for the BP ontology, (iV) the results of GO enrichment analysis for the CC ontology and (v) results of pathway enrichment analysis.

**TABLE S4**. Supplemental file 4 contains the FungiDB enrichment results for the analysis of *A. fumigatus* orthologs of the most-important *Z. tritici* genes. The sheets include (i) BLAST results converting gene identifiers used in this study to *A. fumigatus* identifiers, (ii) the results of GO enrichment analysis for the MF ontology, (iii) the results of GO enrichment analysis for the BP ontology, (iV) the results of GO enrichment analysis for the CC ontology and (v) results of pathway enrichment analysis.

**TABLE S5**. Supplemental file 5 is a large table that provides *log*(*RPKM* + 1) estimates of expression from the Haueisen experiment. Each row represents a gene and columns are samples from the experiment. The column header contains isolate information (Zt05, Zt09 or Zt10), timepoint information (Ta_A, Ta_B, Ta_C or Ta_D) and replicate information (01 or 02).

**TABLE S6**. Supplemental file 6 is a large table that provides *log*(*RPKM* + 1) estimates of expression from the Palma-Guerrero experiment. Each row represents a gene and columns are samples from the experiment. The column header is in the format isolate-replicate-timepoint. Note, that replicate numbers vary as the experimental design of the Palma-Guerrero study used more than 3 replicates for infection assays but only selected the 3 replicates that yielded the best quality RNA for sequencing.

**TABLE S7**. Supplemental file 7 provides information on the development of necrosis symptoms for each isolate including the first appearance of necrosis (measured by days post infection) and the classification of each isolate as early or late onset.

**TABLE S8**. Supplemental file 8 contains the FungiDB enrichment results for the analysis of the most-important *Z. tritici* genes. The sheets include (i) BLAST results converting gene identifiers used in this study to IPO323 identifiers used by FungiDB, (ii) the results of GO enrichment analysis for the BP ontology, (iii) the results of GO enrichment analysis for the MF ontology and (iv) results of pathway enrichment analysis.

**Figure S1.**
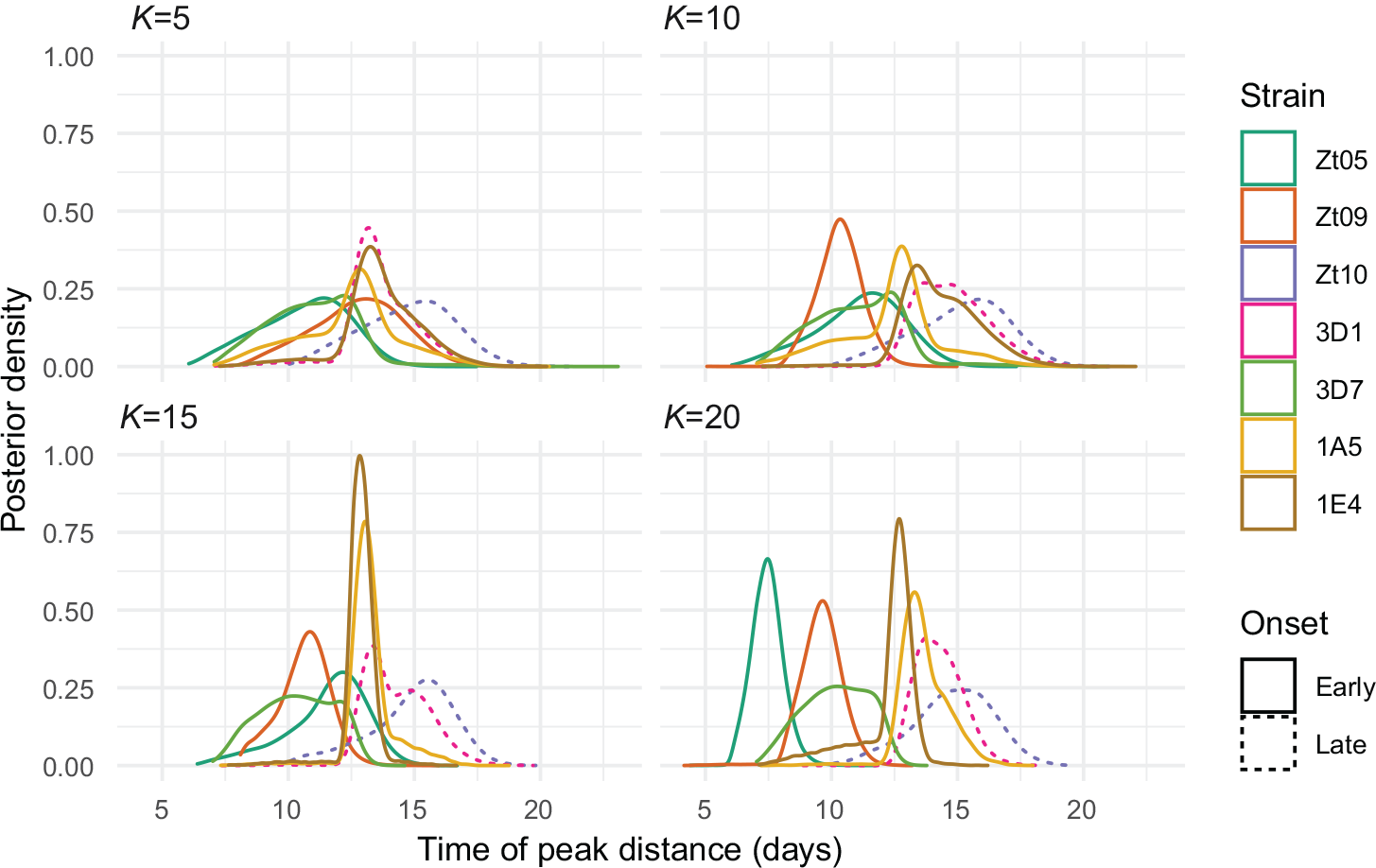
Posterior density of the timing of the third knot, which represents the peak distance relative to the infection start, for each isolate. Results are shown for different values of *K*: 5, 10, 15 and 20. Dashed lines emphasise the two late onset isolates Zt10 (purple) and 3D1 (pink). The peaks for both late onset peaks appear delayed with *K* = 10, 15 and 20 and for isolate Zt10 with *K* = 5.

**Figure S2.**
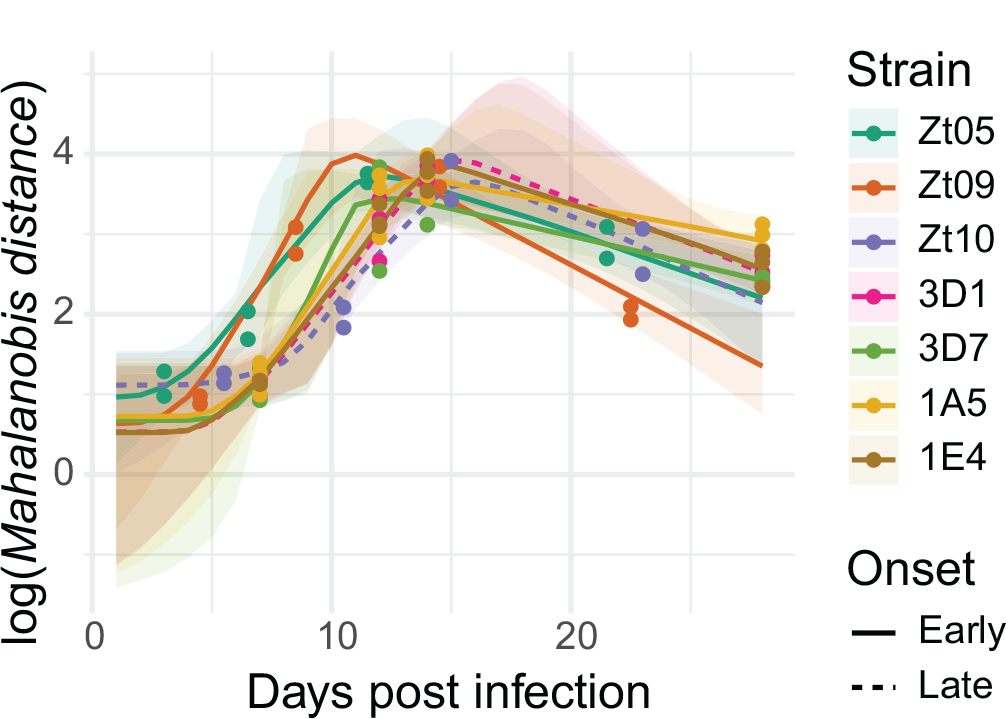
Posterior median expected log-Mahalanobis distance with 95% uncertainty intervals, from the model with data from all isolates at all three stages and using the *K* = 10 most important genes. Different colours represent the isolates and the dashed lines emphasise the two late onset isolates Zt10 (purple) and 3D1 (pink).

**Figure S3.**
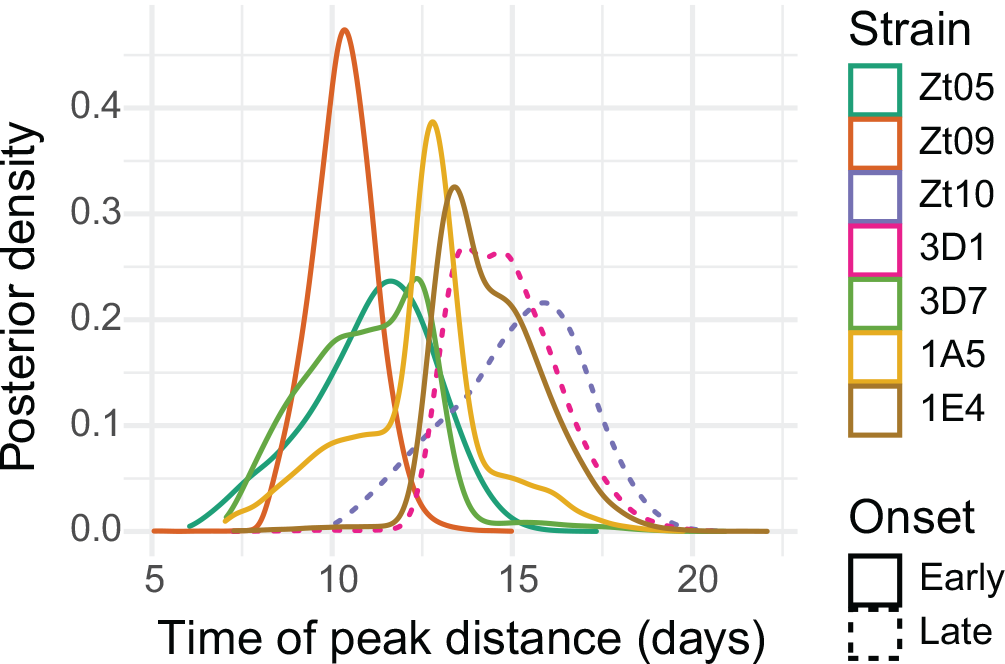
Posterior density for each isolate of the timing of the third knot, which represents the peak distance relative to the infection start, using only the *K* = 10 most important genes. Dashed lines emphasise the two late onset isolates Zt10 (purple) and 3D1 (pink).

## Footnotes

1 Note that, for values of *K* which exceed the number of observations at the initial times (18 in this case), the empirical covariance matrix **Σ** is numerically singular. To avoid this issue, we use the pseudo-inverse in place of **Σ**^*−*1^.

## Notes

### Competing Interest Statement

The authors have declared no competing interest.

## Bibliography

1. S. V. Avery, I. Singleton, N. Magan, and G. H. Goldman. The fungal threat to global food security. Fungal Biology, 123(8):555–557, 2019. ISSN 18786146. doi: 10.1016/j.funbio.2019.03.006. URL https://doi.org/10.1016/j.funbio.2019.03.006.

2. H. N. Fones, D. P. Bebber, T. M. Chaloner, W. T. Kay, G. Steinberg, and S. J. Gurr. Threats to global food security from emerging fungal and oomycete crop pathogens. Nature Food, 1(6):332–342, 2020. ISSN 2662-1355. doi: 10.1038/s43016-020-0075-0. URL http://dx.doi.org/10.1038/s43016-020-0075-0.

3. J. Jones and J. Dangl. The plant immune system. Nature, 444(7117):323–329, 2006. URL https://www.nature.com/articles/nature05286.

4. P. N. Dodds and J. P. Rathjen. Plant immunity: Towards an integrated view of plant-pathogen interactions. Nature Reviews Genetics, 11(8):539–548, 2010. ISSN 14710056. doi: 10.1038/nrg2812. URL https://www.nature.com/articles/nrg2812.

5. L. Lo Presti, D. Lanver, G. Schweizer, S. Tanaka, L. Liang, M. Tollot, A. Zuccaro, S. Reissmann, and R. Kahmann. Fungal Effectors and Plant Susceptibility. Annu. Rev. Plant Biol., 66:513–545, 2015. doi: 10.1146/annurev-arplant-043014-114623. URL https://pubmed.ncbi.nlm.nih.gov/25923844/.

6. S. H. Spoel and X. Dong. How do plants achieve immunity? Defence without specialized immune cells. Nature Reviews Immunology, 12(2):89–100, 2012. ISSN 14741733. doi: 10.1038/nri3141. URL https://www.nature.com/articles/nri3141.

7. H. H. Flor. Current status of the gene-for-gene concept. Annual Review of Phytopathology, 9:275–296, 1971. doi: 10.1146/annurev.py.09.090171.001423. URL https://www.annualreviews.org/doi/abs/10.1146/annurev.py.09.090171.001423.

8. K. E. Hammond-Kosack and J. D. Jones. Resistance gene-dependent plant defense responses. Plant Cell, 8(10):1773–1791, 1996. ISSN 10404651. doi: 10.1105/tpc.8.10.1773. URL https://pubmed.ncbi.nlm.nih.gov/8914325/.

9. A. O’Driscoll, S. Kildea, F. Doohan, J. Spink, and E. Mullins. The wheat-Septoria conflict: A new front opening up? Trends in Plant Science, 19(9):602–610, 2014. ISSN 13601385. doi: 10.1016/j.tplants.2014.04.011. URL http://dx.doi.org/10.1016/j.tplants.2014.04.011.

10. H. Fones and S. Gurr. The impact of Septoria tritici Blotch disease on wheat: An EU perspective. Fungal Genetics and Biology, 79:3–7, 2015. ISSN 10871845. doi: 10.1016/j.fgb.2015.04.004. URL http://linkinghub.elsevier.com/retrieve/pii/S1087184515000705.

11. A. Ponomarenko, S. B. Goodwin, and G. H. J. Kema. Septoria tritici blotch (STB) of wheat. Plant Health Instructor, 2011. doi: 10.1094/PHI-I-2011-0407-01. URL https://doi.org/10.1094/PHI-I-2011-0407-01.

12. S. F. Torriani, J. P. Melichar, C. Mills, N. Pain, H. Sierotzki, and M. Courbot. Zymoseptoria tritici : A major threat to wheat production, integrated approaches to control. Fungal Genetics and Biology, 79:8–12, 2015. ISSN 10871845. doi: 10.1016/j.fgb.2015.04.010. URL http://linkinghub.elsevier.com/retrieve/pii/S1087184515000766.

13. D. Croll, M. H. Lendenmann, E. Stewart, and B. A. McDonald. The Impact of Recombination Hotspots on Genome Evolution of a Fungal Plant Pathogen. Genetics, 201(3): 1213–1228, 2015. ISSN 19432631. doi: 10.1534/genetics.115.180968. URL https://pubmed.ncbi.nlm.nih.gov/26392286/.

14. M. Möller, M. Habig, M. Freitag, and E. H. Stukenbrock. Extraordinary Genome Instability and Widespread Chromosome Rearrangements During Vegetative Growth. Genetics Society of America, 210(October):517–529, 2018. doi: 10.1534/genetics.118.301050. URL https://academic.oup.com/genetics/article/210/2/517/5931050.

15. E. H. Stukenbrock, T. Bataillon, J. Y. Dutheil, T. T. Hansen, R. Li, M. Zala, B. a. McDonald, J. Wang, and M. H. Schierup. The making of a new pathogen: Insights from comparative population genomics of the domesticated wheat pathogen Mycosphaerella graminicola and its wild sister species. Genome Research, 21: 2157–2166, 2011. ISSN 10889051. doi: 10.1101/gr.118851.110. URL https://www.ncbi.nlm.nih.gov/pmc/articles/PMC3227104/.

16. J. Zhan, R. E. Pettway, and B. A. McDonald. The global genetic structure of the wheat pathogen Mycosphaerella graminicola is characterized by high nuclear diversity, low mitochondrial diversity, regular recombination, and gene flow. Fungal Genetics and Biology, 38(3):286–297, 2003. ISSN 10871845. doi: 10.1016/S1087-1845(02)00538-8. URL https://pubmed.ncbi.nlm.nih.gov/12684018/.

17. H. J. Cools and B. A. Fraaije. Are azole fungicides losing ground against Septoria wheat disease? Resistance mechanisms in Mycosphaerella graminicola. Pest management science, 64:681–684, 2008. ISSN 20786913. doi: 10.1002/ps.1568. URL https://pubmed.ncbi.nlm.nih.gov/18366065/.

18. S. F. Torriani, P. C. Brunner, B. a. McDonald, and H. Sierotzki. QoI resistance emerged independently at least 4 times in European populations of Mycosphaerella graminicola. Pest Management Science, 65(2):155–162, 2009. ISSN 1526498X. doi: 10.1002/ps.1662. URL http://doi.wiley.com/10.1002/ps.1662.

19. L. K. Estep, S. F. F. Torriani, M. Zala, N. P. Anderson, M. D. Flowers, B. A. Mcdonald, C. C. Mundt, and P. C. Brunner. Emergence and early evolution of fungicide resistance in North American populations of Zymoseptoria tritici. Plant Pathology, 64(4), 2015. ISSN 13653059. doi: 10.1111/ppa.12314. URL https://bsppjournals.onlinelibrary.wiley.com/doi/10.1111/ppa.12314.

20. J. A. Lucas, N. J. Hawkins, and B. A. Fraaije. The Evolution of Fungicide Resistance. Advances in Applied Microbiology, 90:29–92, 2015. ISSN 00652164. doi: 10.1016/bs.aambs.2014.09.001. URL http://dx.doi.org/10.1016/bs.aambs.2014.09.001.

21. J. J. Blake, P. Gosling, B. A. Fraaije, F. J. Burnett, S. M. Knight, S. Kildea, and N. D. Paveley. Changes in field doseresponse curves for demethylation inhibitor (DMI) and quinone outside inhibitor (QoI) fungicides against Zymoseptoria tritici, related to laboratory sensitivity phenotyping and genotyping assays. Pest Management Science, 74(2):302–313, 2018. ISSN 15264998. doi: 10.1002/ps.4725. URL https://onlinelibrary.wiley.com/doi/abs/10.1002/ps.4725.

22. M. Garnault, C. Duplaix, P. Leroux, G. Couleaud, O. David, A. S. Walker, and F. Carpentier. Large-scale study validates that regional fungicide applications are major determinants of resistance evolution in the wheat pathogen Zymoseptoria tritici in France. New Phytologist, 2020. ISSN 14698137. doi: 10.1111/nph.17107. URL https://nph.onlinelibrary.wiley.com/doi/full/10.1111/nph.17107.

23. F. E. Hartmann, T. Vonlanthen, N. K. Singh, M. C. McDonald, A. Milgate, and D. Croll. The complex genomic basis of rapid convergent adaptation to pesticides across continents in a fungal plant pathogen. Molecular Ecology, (November):1–16, 2020. ISSN 1365294X. doi: 10.1111/mec.15737. URL https://pubmed.ncbi.nlm.nih.gov/33211369/.

24. C. Cowger, M. E. Hoffer, and C. C. Mundt. Specific adaptation by Mycosphaerella graminicola to a resistant wheat cultivar. Plant Pathology, 49 (4):445–451, 2000. ISSN 00320862. doi: 10.1046/j.1365-3059.2000.00472.x. URL https://bsppjournals.onlinelibrary.wiley.com/doi/full/10.1046/j.1365-3059.2000.00472.x.

25. S. B. Goodwin, S. B. M’Barek, B. Dhillon, A. H. J. Wittenberg, C. F. Crane, J. K. Hane, J. Foster, T. a. J. van der Lee, J. Grimwood, A. Aerts, J. Antoniw, A. Bailey, B. Bluhm, J. Bowler, J. Bristow, A. van der Burgt, B. Canto-Canché, A. C. L. Churchill,L. Conde-Ferràez, H. J. Cools, P. M. Coutinho, M. Csukai, P. Dehal, P. de Wit, B. Donzelli, H. C. van de Geest, R. C. H. J. van Ham, K. E. Hammond-Kosack, B. Henrissat, A. Kilian, A. K. Kobayashi, E. Koopmann, Y. Kourmpetis, A. Kuzniar, E. Lindquist, V. Lombard, C. Maliepaard, N. Martins, R. Mehrabi, J. P. H. Nap, A. Ponomarenko, J. J. Rudd, A. Salamov, J. Schmutz, H. J. Schouten, H. Shapiro, I. Stergiopoulos, S. F. F. Torriani, H. Tu, R. P. de Vries, C. Waalwijk, S. B. Ware, A. Wiebenga, L. H. Zwiers, R. P. Oliver, I. V. Grigoriev, and G. H. J. Kema. Finished genome of the fungal wheat pathogen Mycosphaerella graminicola reveals dispensome structure, chromosome plasticity, and stealth pathogenesis. PLoS Genetics, 7(6), 2011. ISSN 15537390. doi: 10.1371/journal.pgen.1002070. URL https://journals.plos.org/plosgenetics/article?id=10.1371/journal.pgen.1002070.

26. T. Badet, U. Oggenfuss, L. Abraham, B. A. McDonald, and D. Croll. A 19-isolate reference-quality global pangenome for the fungal wheat pathogen Zymoseptoria tritici . BMC Biology, (18:20), 2020. doi: 10.1101/803098. URL https://pubmed.ncbi.nlm.nih.gov/32046716/.

27. C. Plissonneau, F. E. Hartmann, and D. Croll. Pangenome analyses of the wheat pathogen Zymoseptoria tritici reveal the structural basis of a highly plastic eukaryotic genome. BMC Biology, 16(1):1–16, 2018. ISSN 17417007. doi: 10.1186/s12915-017-0457-4. URL https://pubmed.ncbi.nlm.nih.gov/29325559/.

28. C. G. McCarthy and D. A. Fitzpatrick. Pan-genome analyses of model fungal species. Microbial Genomics, 5(2), 2019. ISSN 20575858. doi: 10.1099/mgen.0.000243. URL https://www.microbiologyresearch.org/content/journal/mgen/10.1099/mgen.0.000243.

29. M. Habig, J. Quade, and E. H. Stukenbrock. Forward Genetics Approach Reveals Host Genotype-Dependent Importance of Accessory Chromosomes in the Fungal Wheat Pathogen Zymoseptoria tritici . mBio, 8(6):1–16, 2017. ISSN 21507511. doi: 10.1128/mBio.01919-17. URL https://pubmed.ncbi.nlm.nih.gov/29184021/.

30. J. Haueisen, M. Möller, C. J. Eschenbrenner, J. Grandaubert, H. Seybold, H. Adamiak, and E. H. Stukenbrock. Highly flexible infection programs in a specialized wheat pathogen. Ecology and evolution, 9(1):275–294, 2019. doi: 10.1002/ece3.4724. URL https://onlinelibrary.wiley.com/doi/full/10.1002/ece3.4724.

31. J. Palma-Guerrero, X. Ma, S. F. Torriani, M. Zala, C. S. Francisco, F. E. Hartmann, D. Croll, and B. A. McDonald. Comparative transcriptome analyses in Zymoseptoria tritici reveal significant differences in gene expression among strains during plant infection. Molecular Plant-Microbe Interactions, 30(3):231–244, 2017. doi: 10.1094/MPMI-07-16-0146-R. URL https://apsjournals.apsnet.org/doi/10.1094/MPMI-07-16-0146-R.

32. A. Rahman, F. Doohan, and E. Mullins. Quantification of In Planta Zymoseptoria tritici Progression Through Different Infection Phases and Related Association with Components of Aggressiveness. Phytopathology, 110(6):1208–1215, 2020. ISSN 19437684. doi: 10.1094/PHYTO-09-19-0339-R. URL https://pubmed.ncbi.nlm.nih.gov/32133920/.

33. E. S. Luttrell. Parasitism of Fungi on Vascular Plants. Mycologia, 66(1):1–15, 1974. doi: 10.2307/3758447. URL https://www.jstor. org/stable/3758447.

34. Z. Eyal, A. Scharen, J. Prescott, and M. van Ginkel. The Septoria Diseases of Wheat: Concepts and methods of disease management. Mexico, D.F.: CIMMYT, 1987. ISBN 9686127062.

35. G. Kema, D. Yu, F. Rijkenberg, M. Shaw, and R. Baayen. Histology of pathogenesis of Mycosphaerella graminicola in wheat. Phytopathology, 86(7):777–786, 1996. ISSN 0031949X. doi: 10.1094/Phyto-86-777.

36. R. Mehrabi, T. van der Lee, C. Waalwijk, and G. H. J. Kema. MgSlt2, a Cellular Integrity MAP Kinase Gene of the Fungal Wheat Pathogen Mycosphaerella graminicola, Is Dispensable for Penetration but Essential for Invasive Growth. Molecular Plant-Microbe Interactions, 19(4):389–398, 2006. ISSN 0894-0282. doi: 10.1094/MPMI-19-0389. URL http://apsjournals.apsnet.org/doi/abs/10.1094/MPMI-19-0389.

37. N. Shetty, B. Kristensen, M.-A. Newman, K. Møller, P. Gregersen, and H. Jørgensen. Association of hydrogen peroxide with restriction of Septoria tritici in resistant wheat. Physiological and Molecular Plant Pathology, 62(6):333–346, 2003. ISSN 08855765. doi: 10.1016/S0885-5765(03)00079-1. URL http://linkinghub.elsevier.com/retrieve/pii/S0885576503000791.

38. N. P. Shetty, R. Mehrabi, H. Lütken, A. Haldrup, G. H. J. Kema, D. B. Collinge, and H. J. L. Jørgensen. Role of hydrogen peroxide during the interaction between the hemibiotrophic fungal pathogen Septoria tritici and wheat. New Phytologist, 174 (3):637–647, 2007. ISSN 0028646X. doi: 10.1111/j.1469-8137.2007.02026.x. URL https://nph.onlinelibrary.wiley.com/doi/10.1111/j.1469-8137.2007.02026.x.

39. J. Keon, J. Antoniw, R. Carzaniga, S. Deller, J. L. Ward, J. M. Baker, M. H. Beale, K. Hammond-Kosack, and J. J. Rudd. Transcriptional adaptation of Mycosphaerella graminicola to programmed cell death (PCD) of its susceptible wheat host. Molecular plant-microbe interactions : MPMI, 20(2):178–193, 2007. ISSN 0894-0282. doi: 10.1094/MPMI-20-2-0178. URL https://apsjournals.apsnet.org/doi/10.1094/MPMI-20-2-0178.

40. G. H. J. Kema, T. A. J. van der Lee, O. Mendes, E. C. P. Verstappen, R. K. Lankhorst, H. Sandbrink, A. van der Burgt, L.-H. Zwiers, M. Csukai, and C. Waalwijk. Large-Scale Gene Discovery in the Septoria Tritici Blotch Fungus Mycosphaerella graminicola with a Focus on In Planta Expression. Molecular plant-microbe interactions : MPMI, 21 (9):1249–60, 2008. ISSN 0894-0282. doi: 10.1094/MPMI-21-9-1249. URL http://apsjournals.apsnet.org/doi/abs/10.1094/MPMI-21-9-1249.

41. J. J. Rudd, J. Keon, and K. E. Hammond-Kosack. The wheat mitogen-activated protein kinases TaMPK3 and TaMPK6 are differentially regulated at multiple levels during compatible disease interactions with Mycosphaerella graminicola. Plant Physiology, 147(2):802–815, 2008. ISSN 00320889. doi: 10.1104/pp.108.119511. URL https://pubmed.ncbi.nlm.nih.gov/18441220/.

42. R. Marshall, A. Kombrink, J. Motteram, E. Loza-Reyes, J. Lucas, K. E. Hammond-Kosack, B. P. H. J. Thomma, and J. J. Rudd. Analysis of Two in Planta Expressed LysM Effector Homologs from the Fungus Mycosphaerella graminicola Reveals Novel Functional Properties and Varying Contributions to Virulence on Wheat. Plant Physiology, 156(2):756–769, 2011. ISSN 0032-0889. doi: 10.1104/pp.111.176347. URL https://academic.oup.com/plphys/article/156/2/756/6108806.

43. A. Sánchez-Vallet, M. C. Mcdonald, P. S. Solomon, and B. a. Mcdonald. Is Zy-moseptoria tritici a hemibiotroph? Fungal Genetics and Biology, 79:29–32, 2015. doi: 10.1016/j.fgb.2015.04.001. URL https://pubmed.ncbi.nlm.nih.gov/26092787/.

44. J. Rudd, K. Kanyuka, K. Hassani-Pak, M. Derbyshire, A. Andongabo, J. Devonshire Lysenko, M. Saqi, N. Desai, S. Powers, J. Hooper, L. Ambroso, A. Bharti, A. Farmer, K. Hammond-kosack, R. Dietrich, and M. Courbot. Transcriptome and metabolite profiling the infection cycle of Zymoseptoria tritici on wheat (Triticum aestivum) reveals a biphasic interaction with plant immunity involving differential pathogen chromosomal contributions, and a variation on the hemibiotrophic lifestyle definition. Plant Physiology, 167(March):pp.114.255927, 2015. ISSN 0032-0889. doi: 10.1104/pp.114.255927. URL https://pubmed.ncbi.nlm.nih.gov/25596183/.

45. E. Fantozzi, S. Kilaru, S. J. Gurr, and G. Steinberg. Asynchronous development of Zymoseptoria tritici infection in wheat. Fungal Genetics and Biology, 146(August 2020):103504, 2021. ISSN 10960937. doi: 10.1016/j.fgb.2020.103504. URL https://pubmed.ncbi.nlm.nih.gov/33326850/.

46. K. E. Duncan and R. J. Howard. Cytological analysis of wheat infection by the leaf blotch pathogen Mycosphaerella graminicola. Mycological Research, 104(9):1074–1082, 2000. ISSN 09537562. doi: 10.1017/S0953756299002294. URL https://www.sciencedirect.com/science/article/pii/S0953756208617591.

47. S. Pnini-Cohen, A. Zilberstein, S. Schuster, A. Sharon, and Z. Eyal. Elucidation of Septoria tritici x Wheat Interactions Using GUS-Expressing Isolates. Phytopathology, 90(3):297–304, 2000. ISSN 0031949X. doi: 10.1094/PHYTO.2000.90.3.297. URL https://pubmed.ncbi.nlm.nih.gov/18944623/.

48. S. Deller, K. E. Hammond-Kosack, and J. J. Rudd. The complex interactions between host immunity and non-biotrophic fungal pathogens of wheat leaves. Journal of Plant Physiology, 168(1):63–71, 2011. ISSN 01761617. doi: 10.1016/j.jplph.2010.05.024. URL https://pubmed.ncbi.nlm.nih.gov/20688416/.

49. R. Dean, J. A. L. Van Kan, Z. A. Pretorius, K. E. Hammond-Kosack, A. Di Pietro, P. D. Spanu, J. J. Rudd, M. Dickman, R. Kahmann, J. Ellis, and G. D. Foster. The Top 10 fungal pathogens in molecular plant pathology. Molecular Plant Pathology, 13 (4):414–430, 2012. ISSN 14646722. doi: 10.1111/j.1364-3703.2011.00783.x. URL https://pubmed.ncbi.nlm.nih.gov/22471698/.

50. G. Steinberg. Cell biology of Zymoseptoria tritici: Pathogen cell organization and wheat infection. Fungal Genetics and Biology, 79:17–23, 2015. ISSN 10871845. doi: 10.1016/j.fgb.2015.04.002. URL http://linkinghub.elsevier.com/retrieve/pii/S108718451500064X.

51. W. E. Sackston. A Possible Mechanism Of Dispersal Of Septoria Spores. Canadian Journal Of Plant Science, 50(2):155–157, 1970. doi: 10.4141/cjps70-029. URL https://cdnsciencepub.com/doi/10.4141/cjps70-029.

52. H. N. Fones, C. J. Eyles, W. Kay, J. Cowper, and S. J. Gurr. A role for random, humidity-dependent epiphytic growth prior to invasion of wheat by Zymoseptoria tritici. Fungal Genetics and Biology, 106(May):51–60, 2017. ISSN 10960937. doi: 10.1016/j.fgb.2017.07.002. URL http://dx.doi.org/10.1016/j.fgb.2017.07.002.

53. W.-s. Lee, J. J. Rudd, K. E. Hammond-kosack, and K. K. Kanyuka. Mycosphaerella graminicola LysM effector-mediated stealth pathogenesis subverts recognition through both CERK1 and CEBiP homologues in wheat. Molecular plantmicrobe interactions : MPMI, 27(3):236–243, 2013. ISSN 0894-0282. doi: 10.1094/MPMI-07-13-0201-R. URL http://www.ncbi.nlm.nih.gov/pubmed/24073880.

54. A. Mirzadi Gohari, S. B. Ware, A. H. J. Wittenberg, R. Mehrabi, S. Ben M’Barek, E. C. P. Verstappen, T. A. J. van der Lee, O. Robert, H. J. Schouten, P. P. J. G. M. de Wit, and G. H. J. Kema. Effector discovery in the fungal wheat pathogen Zymoseptoria tritici. Molecular plant pathology, (July), 2015. ISSN 1364-3703. doi: 10.1111/mpp.12251. URL http://www.ncbi.nlm.nih.gov/pubmed/25727413.

55. S. Poppe, L. Dorsheimer, P. Happel, and E. H. Stukenbrock. Rapidly Evolving Genes Are Key Players in Host Specialization and Virulence of the Fungal Wheat Pathogen Zymoseptoria tritici (Mycosphaerella graminicola). PLOS Pathogens, 11 (7):e1005055, 2015. ISSN 1553-7374. doi: 10.1371/journal.ppat.1005055. URL http://dx.plos.org/10.1371/journal.ppat.1005055.

56. G. J. Kettles, C. Bayon, G. Canning, J. J. Rudd, and K. Kanyuka. Apoplastic recognition of multiple candidate effectors from the wheat pathogen Zymoseptoria tritici in the nonhost plant Nicotiana benthamiana. New Phytologist, (213):338–350, 2017. doi: 10.1111/nph.14215. URL https://pubmed.ncbi.nlm.nih.gov/27696417/.

57. M. C. McDonald, L. McGinness, J. K. Hane, A. H. Williams, A. Milgate, and P. S. Solomon. Utilizing Gene Tree Variation to Identify Candidate Effector Genes in Zymoseptoria tritici. Genes Genomes Genetics, 6(4):779–791, 2016. ISSN 2160-1836. doi: 10.1534/g3.115.025197. URL https://pubmed.ncbi.nlm.nih.gov/26837952/.

58. R. King, M. Urban, R. P. Lauder, N. Hawkins, M. Evans, A. Plummer, K. Halsey, A. Lovegrove, K. Hammond-Kosack, and J. J. Rudd. A conserved fungal glycosyltransferase facilitates pathogenesis of plants by enabling hyphal growth on solid surfaces. PLoS Pathogens, 13(10):1–26, 2017. ISSN 1553-7374. doi: 10.1371/journal.ppat.1006672. URL http://journals.plos.org/plospathogens/article/file?id=10.1371/journal.ppat.1006672&type=printable.

59. C. Plissonneau, J. Benevenuto, N. Mohd-Assaad, S. Fouché, F. E. Hartmann, and D. Croll. Using Population and Comparative Genomics to Understand the Genetic Basis of Effector-Driven Fungal Pathogen Evolution. Frontiers in Plant Science, 8: 1–15, 2017. ISSN 1664-462X. doi: 10.3389/fpls.2017.00119. URL https://pubmed.ncbi.nlm.nih.gov/28217138/.

60. Z. Zhong, T. C. Marcel, F. E. Hartmann, X. Ma . Plissonneau, M. Zala, A. Ducasse, J. Confais, J. Compain, N. Lapalu, J. Amselem, B. A. McDonald, D. Croll, and J. Palma-Guerrero. A small secreted protein in Zymoseptoria tritici is responsible for avirulence on wheat cultivars carrying the Stb6 resistance gene. New Phytologist, 214(2), 2017. ISSN 14698137. doi: 10.1111/nph.14434. URL https://nph.onlinelibrary.wiley.com/doi/full/10.1111/nph.14434.

61. S. Fouché, C. Plissonneau, and D. Croll. The birth and death of effectors in rapidly evolving filamentous pathogen genomes. Current Opinion in Microbiology, 46:34–42, 2018. ISSN 18790364. doi: 10.1016/j.mib.2018.01.020. URL https://pubmed.ncbi.nlm.nih.gov/29455143/.

62. G. H. Kema, A. Mirzadi Gohari, L. Aouini, H. A. Gibriel, S. B. Ware, F. van Den Bosch, R. Manning-Smith, V. Alonso-Chavez, J. Helps, S. Ben M’Barek, R. Mehrabi, C. Diaz-Trujillo, E. Zamani, H. J. Schouten, T. A. van der Lee, C. Waalwijk, M. A. de Waard, P. J. de Wit, E. C. Verstappen, B. P. Thomma, H. J. Meijer, and M. F. Seidl. Stress and sexual reproduction affect the dynamics of the wheat pathogen effector AvrStb6 and strobilurin resistance. Nature Genetics, 50(March):1–6, 2018. ISSN 15461718. doi: 10.1038/s41588-018-0052-9. URL http://dx.doi.org/10.1038/s41588-018-0052-9.

63. G. J. Kettles, C. Bayon, C. A. Sparks, G. Canning, K. Kanyuka, and J. J. Rudd. Characterization of an antimicrobial and phytotoxic ribonuclease secreted by the fungal wheat pathogen Zymoseptoria tritici . New Phytologist, 217(1):320–331, 2018. ISSN 14698137. doi: 10.1111/nph.14786. URL https://nph.onlinelibrary.wiley.com/doi/full/10.1111/nph.14786.

64. L. Meile, D. Croll, P. C. Brunner, C. Plissonneau, F. E. Hartmann, B. A. McDonald, and A. Sánchez-Vallet. A fungal avirulence factor encoded in a highly plastic genomic region triggers partial resistance to septoria tritici blotch. The New Phytologist, 219 (3):1048–1061, 2018. doi: 10.1111/nph.15180. URL https://www.ncbi.nlm.nih.gov/pmc/articles/PMC6055703/.

65. C. Saintenac, W.-S. Lee, F. Cambon, J. J. Rudd, R. C. King, W. Marande, S. J. Powers, H. Bergès, A. L. Phillips, C. Uauy, K. E. Hammond-Kosack, T. Langin, and K. Kanyuka. Wheat receptor-kinase-like protein Stb6 controls gene-for-gene resistance to fungal pathogen Zymoseptoria tritici . Nature Genetics, 2018. ISSN 1061-4036. doi: 10.1038/s41588-018-0051-x. URL http://www.nature.com/articles/s41588-018-0051-x.

66. S. J. Karki, A. Reilly, B. Zhou, M. Mascarello, J. Burke, F. Doohan, D. Douchkov, P. Schweizer, and A. Feechan. A small secreted protein from Zymoseptoria tritici interacts with a wheat E3 ubiquitin ligase to promote disease. Journal of Experimental Botany, 72(2):733–746, 2020. ISSN 0022-0957. doi: 10.1093/jxb/eraa489. URL https://pubmed.ncbi.nlm.nih.gov/33095257/.

67. A. Sánchez-Vallet, H. Tian, L. Rodriguez-Moreno, D. J. Valkenburg, R. Saleem-Batcha, S. Wawra, A. Kombrink, L. Verhage, R. de Jonge, H. P. van Esse, A. Zuccaro, D. Croll, J. R. Mesters, and B. P. Thomma. A secreted LysM effector protects fungal hyphae through chitin-dependent homodimer polymerization. PLoS Pathogens, 16(6 June):1–21, 2020. ISSN 15537374. doi: 10.1371/journal.ppat.1008652. URL https://journals.plos.org/plospathogens/article?id=10.1371/journal.ppat.1008652.

68. C. Stephens, F. Ölmez, H. Blyth, M. McDonald, A. Bansal, E. Burcu Turgay, F. Hahn, C. Saintenac, V. Nekrasov, P. Solomon, A. Milgate, B. Fraaije, J. Rudd, and K. Kanyuka. Remarkable recent changes in genetic diversity of the avirulence gene AvrStb6 in global populations of the wheat pathogen Zymoseptoria tritici . Molecular Plant Pathology, page 2020.09.18.303370, 2021. doi: 10.1111/mpp.13101. URL https://bsppjournals.onlinelibrary.wiley.com/doi/full/10.1111/mpp.13101.

69. A. Cousin, R. Mehrabi, M. Guilleroux, M. Dufresne, T. Van Der Lee, C. Waalwijk, T. Langin, and G. H. J. Kema. The MAP kinase-encoding gene MgFus3 of the non-appressorium phytopathogen Mycosphaerella graminicola is required for penetration and in vitro pycnidia formation. Molecular Plant Pathology, 7(4):269–278, 2006. ISSN 14646722. doi: 10.1111/j.1364-3703.2006.00337.x. URL https://pubmed.ncbi.nlm.nih.gov/20507446/.

70. R. Mehrabi, L.-H. Zwiers, M. a. de Waard, and G. H. J. Kema. MgHog1 regulates dimorphism and pathogenicity in the fungal wheat pathogen Mycosphaerella graminicola. Molecular plant-microbe interactions : MPMI, 19(11):1262–1269, 2006. ISSN 0894-0282. doi: 10.1094/MPMI-19-1262. URL https://apsjournals.apsnet.org/doi/10.1094/MPMI-19-1262.

71. A. Mirzadi Gohari, R. Mehrabi, O. Robert, I. A. Ince, S. Boeren, M. Schuster, G. Steinberg, P. J. G. M. de Wit, and G. H. J. Kema. Molecular characterization and functional analyses of ZtWor1,a transcriptional regulator of the fungal wheat pathogen Zymoseptoria tritici . Molecular Plant Pathology, 15(4), 2014. doi: 10.1111/mpp.12102.URL https://pubmed.ncbi.nlm.nih.gov/24341593/.

72. N. Mohammadi, R. Mehrabi, A. Mirzadi Gohari, E. Mohammadi Goltapeh, N. Safaie, and G. H. Kema. The ZtVf1 transcription factor regulates development and virulence in the foliar wheat pathogen Zymoseptoria tritici . Fungal Genetics and Biology, 109(May):26–35, 2017. ISSN 10960937. doi: 10.1016/j.fgb.2017.10.003. URL https://pubmed.ncbi.nlm.nih.gov/29031630/.

73. M. Habig, S. M. Bahena-Garrido, F. Barkmann, J. Haueisen, and E. H. Stukenbrock. The transcription factor Zt107320 affects the dimorphic switch, growth and virulence of the fungal wheat pathogen Zymoseptoria tritici . Molecular Plant Pathology, page 595454, 2019. doi: 10.1111/mpp.12886. URL https://bsppjournals.onlinelibrary.wiley.com/doi/10.1111/mpp.12886.

74. N. Mohammadi, R. Mehrabi, A. Mirzadi Gohari, M. Roostaei, E. Mohammadi Goltapeh, N. Safaie, and G. H. Kema. MADS-Box Transcription Factor ZtRlm1 Is Responsible for Virulence and Development of the Fungal Wheat Pathogen Zymoseptoria tritici. Frontiers in Microbiology, 11(August):1–13, 2020. ISSN 1664302X. doi: 10.3389/fmicb.2020.01976. URL https://www.frontiersin.org/articles/10.3389/fmicb.2020.01976/full.

75. P. C. Brunner, S. F. F. Torriani, D. Croll, E. H. Stukenbrock, and B. a. Mc-Donald. Coevolution and life cycle specialization of plant cell wall degrading enzymes in a hemibiotrophic pathogen. Molecular biology and evolution, 30(6):1337–47, 2013. ISSN 1537-1719. doi: 10.1093/molbev/mst041. URL http://www.pubmedcentral.nih.gov/articlerender.fcgi?artid=3649673&tool=pmcentrez&rendertype=abstract.

76. F. Yang, W. Li, and H. J. L. Jørgensen. Transcriptional reprogramming of wheat and the hemibiotrophic pathogen Septoria tritici during two phases of the compatible interaction. PLoS ONE, 8(11):1–15, 2013. ISSN 19326203. doi: 10.1371/journal.pone.0081606. URL https://journals.plos.org/plosone/article?id=10.1371/journal.pone.0081606.

77. R. Kellner, A. Bhattacharyya, S. Poppe, T. Y. Hsu, R. B. Brem, and E. H. Stukenbrock. Expression Profiling of the Wheat Pathogen Zymoseptoria tritici Reveals Genomic Patterns of Transcription and Host-Specific Regulatory Programs. Genome Biology and Evolution, 6(6):1353–1365, 2014. ISSN 1759-6653. doi: 10.1093/gbe/evu101. URL http://gbe.oxfordjournals.org/cgi/doi/10.1093/gbe/evu101.

78. J. Grandaubert, A. Bhattacharyya, and E. H. Stukenbrock. RNA-seq-based gene annotation and comparative genomics of four fungal grass pathogens in the genus Zymoseptoria identify novel orphan genes and species-specific invasions of transposable elements. G3: Genes, Genomes, Genetics, 5(7):1323–1333, 2015. doi: 10.1534/g3.115.017731. URL https://pubmed.ncbi.nlm.nih.gov/25917918/.

79. J. Palma-Guerrero, S. F. F. Torriani, M. Zala, D. Carter, J. J. Rudd, B. A. Mcdonald, and D. Croll. Comparative transcriptome analyses of Zymoseptoria tritici strains show complex lifestyle transitions and intraspecific variability in transcription profiles. Molecular Plant Pathology, pages 1–41, 2016. ISSN 1364-3703. doi: 10.1111/mpp.12333. URL https://pubmed.ncbi.nlm.nih.gov/26610174/.

80. S. Fouché, T. Badet, U. Oggenfuss, C. Plissonneau, C. S. Francisco, and D. Croll. Stress-Driven Transposable Element De-repression Dynamics and Virulence Evolution in a Fungal Pathogen. Molecular Biology and Evolution, 37(1):221–239, 2019. ISSN 15371719. doi: 10.1093/molbev/msz216. URL https://pubmed.ncbi.nlm.nih.gov/31553475/.

81. L. Meile, J. Peter, G. Puccetti, J. Alassimone, B. A. McDonald, and A. Sánchez-Vallet. Chromatin dynamics contribute to the spatiotemporal expression pattern of virulence genes in a fungal plant pathogen. mBio, 11(5):1–18, 2020. ISSN 21507511. doi: 10.1128/mBio.02343-20. URL https://pubmed.ncbi.nlm.nih.gov/33024042/.

82. J. Sperschneider. Machine learning in plant–pathogen interactions: empowering biological predictions from field scale to genome scale. New Phytologist, 228(1):35–41, 2020. doi: 10.1111/nph.15771. URL https://pubmed.ncbi.nlm.nih.gov/30834534/.

83. R. Shaik and W. Ramakrishna. Machine learning approaches distinguish multiple stress conditions using stress-responsive genes and identify candidate genes for broad resistance in rice. Plant physiology, 164(1):481–495, 2014. doi: 10.1104/pp.113.225862. URL https://pubmed.ncbi.nlm.nih.gov/24235132/.

84. S. Liu, Y. Liu, J. Zhao, S. Cai, H. Qian, K. Zuo, L. Zhao, and L. Zhang. A computational interactome for prioritizing genes associated with complex agronomic traits in rice (Oryza sativa). The Plant Journal, 90(1):177–188, 2017. doi: 10.1111/tpj.13475. URL https://pubmed.ncbi.nlm.nih.gov/28074633/.

85. J. Grandaubert, J. Y. Dutheil, and E. H. Stukenbrock. The genomic determinants of adaptive evolution in a fungal pathogen. Evolution Letters, 3(3):299–312, 2019. doi: 10.1002/evl3.117. URL https://www.ncbi.nlm.nih.gov/pmc/articles/PMC6546377/.

86. X. Chen and H. Ishwaran. Random forests for genomic data analysis. Genomics, 99 (6):323–329, 2012. doi: 10.1016/j.ygeno.2012.04.003. URL https://pubmed.ncbi.nlm.nih.gov/22546560/.

87. R. Díaz-Uriarte and S. A. De Andres. Gene selection and classification of microarray data using random forest. BMC bioinformatics, 7(1):1–13, 2006. doi: 10.1186/1471-2105-7-3. URL https://pubmed.ncbi.nlm.nih.gov/16398926/.

88. M. D. Robinson, D. J. McCarthy, and G. K. Smyth. edgeR: a Bioconductor package for differential expression analysis of digital gene expression data. Bioinformatics, 26(1):139–140, 2010. doi: 10.1093/bioinformatics/btp616. URL https://pubmed.ncbi.nlm.nih.gov/19910308/.

89. M. Ashburner, C. Ball, J. Blake, D. Botstein, H. Butler, J. Cherry, A. Davis, K. Dolinski, S. Dwight, J. Eppig, et al. Gene ontology: tool for the unification of biology. The Gene Ontology Consortium. Nature Genetics, 25(1):25–29, 2000. doi: 10.1038/75556. URL https://pubmed.ncbi.nlm.nih.gov/10802651/.

90. L.-H. Zwiers and M. A. De Waard. Characterization of the ABC transporter genes MgAtr1 and MgAtr2 from the wheat pathogen Mycosphaerella graminicola. Fungal Genetics and Biology, 30(2):115–125, 2000. doi: 10.1006/fgbi.2000.1209. URL https://pubmed.ncbi.nlm.nih.gov/11017767/.

91. I. V. Grigoriev, R. Nikitin, S. Haridas, A. Kuo, R. Ohm, R. Otillar, R. Riley, A. Salamov, X. Zhao, F. Korzeniewski, et al. MycoCosm portal: gearing up for 1000 fungal genomes. Nucleic acids research, 42(D1):D699–D704, 2014. doi: 10.1093/nar/gkt1183. URL https://pubmed.ncbi.nlm.nih.gov/24297253/.

92. E. Y. Basenko, J. A. Pulman, A. Shanmugasundram, O. S. Harb, K. Crouch, D. Starns, S. Warrenfeltz, C. Aurrecoechea, C. J. Stoeckert, J. C. Kissinger, et al. FungiDB: An Integrated Bioinformatic Resource for Fungi and Oomycetes. Journal of Fungi, 4 (1):39, 2018. doi: 10.3390/jof4010039. URL https://pubmed.ncbi.nlm.nih.gov/30152809/.

93. B. Cheifet. Where is genomics going next? Genome Biology, 20(1):NA–NA, 2019. doi: 10.1186/s13059-019-1626-2. URL https://genomebiology.biomedcentral.com/articles/10.1186/s13059-019-1626-2.

94. K. Mochida, S. Koda, K. Inoue, and R. Nishii. Statistical and machine learning approaches to predict gene regulatory networks from transcriptome datasets. Frontiers in Plant Science, 9:1770, 2018. doi: 10.3389/fpls.2018.01770. URL https://www.frontiersin.org/articles/10.3389/fpls.2018.01770/full.

95. L. Guo, G. Zhao, J.-R. Xu, H. C. Kistler, L. Gao, and L.-J. Ma. Compartmentalized gene regulatory network of the pathogenic fungus Fusarium graminearum. The New Phytologist, 211(2):527, 2016. doi: 10.1111/nph.13912. URL https://pubmed.ncbi.nlm.nih.gov/26990214/.

96. L.-H. Zwiers, I. Stergiopoulos, M. M. Gielkens, S. D. Goodall, and M. A. De Waard. ABC transporters of the wheat pathogen Mycosphaerella graminicola function as protectants against biotic and xenobiotic toxic compounds. Molecular Genetics and Genomics, 269(4):499–507, 2003. doi: 10.1007/s00438-003-0855-x. URL https://pubmed.ncbi.nlm.nih.gov/12768412/.

97. I. Stergiopoulos, L.-H. Zwiers, and M. A. De Waard. The ABC transporter MgAtr4 is a virulence factor of Mycosphaerella graminicola that affects colonization of substomatal cavities in wheat leaves. Molecular plant-microbe interactions, 16(8):689–698, 2003. doi: 10.1094/MPMI.2003.16.8.689. URL https://pubmed.ncbi.nlm.nih.gov/12906113/.

98. R. Roohparvar, M. A. De Waard, G. H. Kema, and L.-H. Zwiers. MgMfs1, a major facilitator superfamily transporter from the fungal wheat pathogen Mycosphaerella graminicola, is a strong protectant against natural toxic compounds and fungicides. Fungal Genetics and Biology, 44(5):378–388, 2007. doi: 10.1016/j.fgb.2006.09.007. URL https://pubmed.ncbi.nlm.nih.gov/17107817/.

99. J. Heller and P. Tudzynski. Reactive oxygen species in phytopathogenic fungi: signaling, development, and disease. Annual review of phytopathology, 49:369–390, 2011. doi: 10.1146/annurev-phyto-072910-095355. URL https://pubmed.ncbi.nlm.nih.gov/21568704/.

100. A. J. Brown, K. Haynes, and J. Quinn. Nitrosative and oxidative stress responses in fungal pathogenicity. Current opinion in microbiology, 12(4):384–391, 2009. doi: 10.1016/j.mib.2009.06.007. URL https://pubmed.ncbi.nlm.nih.gov/19616469/.

101. L. J. Heung, C. Luberto, and M. Del Poeta. Role of sphingolipids in microbial pathogenesis. Infection and immunity, 74(1):28–39, 2006. doi: 10.1128/IAI.74.1.28-39.2006.

102. J. Cheng, T.-S. Park, L.-C. Chio, A. S. Fischl, and S. Y. Xiang. Induction of apoptosis by sphingoid long-chain bases in Aspergillus nidulans. Molecular and cellular biology, 23(1):163–177, 2003. doi: 10.1128/MCB.23.1.163-177.2003. URL https://pubmed.ncbi.nlm.nih.gov/12482970/.

103. C. Luberto, D. L. Toffaletti, E. A. Wills, S. C. Tucker, A. Casadevall, J. R. Perfect, Y. A. Hannun, and M. Del Poeta. Roles for inositol-phosphoryl ceramide synthase 1 (IPC1) in pathogenesis of C. neoformans. Genes & development, 15(2):201–212, 2001. doi: 10.1101/gad.856001. URL https://pubmed.ncbi.nlm.nih.gov/11157776/.

104. X. Qiang, Z. Kou, G. Fang, and Y. Wang. Scoring amino acid mutations to predict avian-to-human transmission of avian influenza viruses. Molecules, 23(7):1584, 2018. doi: 10.3390/molecules23071584. URL https://pubmed.ncbi.nlm.nih.gov/29966263/.

105. L. K. Mehra, C. Cowger, K. Gross, and P. S. Ojiambo. Predicting pre-planting risk of Stagonospora nodorum blotch in winter wheat using machine learning models. Frontiers in plant science, 7:390, 2016. doi: 10.3389/fpls.2016.00390. URL https://pubmed.ncbi.nlm.nih.gov/27064542/.

106. M. Pineda, M. L. Pérez-Bueno, and M. Barón. Detection of Bacterial Infection in Melon Plants by Classification Methods Based on Imaging Data. Frontiers in plant science, 9:164, 2018. doi: 10.3389/fpls.2018.00164. URL https://pubmed.ncbi.nlm.nih.gov/29491881/.

107. F. Odilbekov, R. Armoniené, T. Henriksson, and A. Chawade. Proximal Phenotyping and Machine Learning Methods to Identify Septoria Tritici Blotch Disease Symptoms in Wheat. Frontiers in plant science, 9:685, 2018. doi: 10.3389/fpls.2018.00685. URL https://pubmed.ncbi.nlm.nih.gov/29875788/.

108. J. C. Tovar, J. S. Hoyer, A. Lin, A. Tielking, S. T. Callen, S. Elizabeth Castillo, M. Miller, M. Tessman, N. Fahlgren, J. C. Carrington, et al. Raspberry Pi–powered imaging for plant phenotyping. Applications in plant sciences, 6(3):e1031, 2018. doi: 10.1002/aps3.1031. URL https://pubmed.ncbi.nlm.nih.gov/29732261/.

109. B. Mishra, N. Kumar, and M. S. Mukhtar. Systems Biology and Machine Learning in Plant–Pathogen Interactions. Molecular Plant-Microbe Interactions, 32(1):45–55, 2019. doi: 10.1094/MPMI-08-18-0221-FI. URL https://pubmed.ncbi.nlm.nih.gov/30418085/.

110. T. Barrett, S. E. Wilhite, P. Ledoux, C. Evangelista, I. F. Kim, M. Tomashevsky, K. A. Marshall, K. H. Phillippy, P. M. Sherman, M. Holko, et al. NCBI GEO: archive for functional genomics data sets—update. Nucleic acids research, 41(D1):D991–D995, 2013. doi: 10.1093/nar/gks1193. URL https://www.ncbi.nlm.nih.gov/pmc/articles/PMC3531084/.

111. Y. Benjamini and Y. Hochberg. Controlling the false discovery rate: a practical and powerful approach to multiple testing. Journal of the Royal Statistical Society. Series B (Methodological), 57(1):289–300, 1995. doi: 10.1111/j.2517-6161.1995.tb02031.x. URL https://rss.onlinelibrary.wiley.com/doi/10.1111/j.2517-6161.1995.tb02031.x.

112. L. Breiman. Random forests. Machine learning, 45(1):5–32, 2001. doi: 10.1023/A:1010933404324.

113. M. N. Wright and A. Ziegler. ranger: A Fast Implementation of Random Forests for High Dimensional Data in C++ and R. Journal of Statistical Software, 77(1):1–17, 2017. doi: 10.18637/jss.v077.i01. URL https://www.jstatsoft.org/article/view/v077i01.

114. P. de Valpine, D. Turek, C. J. Paciorek, C. Anderson-Bergman, D. T. Lang, and R. Bodik. Programming with models: writing statistical algorithms for general model structures with NIMBLE. Journal of Computational and Graphical Statistics, 26(2): 403–413, 2017. doi: 10.1080/10618600.2016.1172487. URL https://www.tandfonline.com/doi/abs/10.1080/10618600.2016.1172487.

